# Dispersal and polyandry evolve as competing inbreeding avoidance strategies

**DOI:** 10.1101/2021.10.10.463818

**Authors:** Greta Bocedi

## Abstract

Understanding evolution of complex life-histories requires explicitly considering their multiple interactions, feedbacks, and shared drivers. Inbreeding depression is hypothesized to drive evolution of two life-histories which have far-reaching ecological and evolutionary consequence: dispersal and polyandry. Yet, the role of inbreeding depression in the separate evolution of these key life-histories is still debated, while the possibility for their joint evolution and consequent covariation has not been considered. I propose that dispersal and polyandry might be competing means of inbreeding avoidance which negatively feedback on each other’s evolution. Using a genetically explicit individual-based model, I first demonstrate that inbreeding depression can drive the separate evolution of dispersal and polyandry. Although this is largely known for dispersal, it is not as well established for polyandry evolution, which generally remains an evolutionary puzzle. Here, I show that polyandry can indeed evolve as means of indirect inbreeding avoidance in spatially structured populations. Second, when dispersal and polyandry can evolve jointly, a negative feedback emerges, such that they evolve as alternative inbreeding avoidance strategies across replicate systems, especially if there are fitness costs associated. Finally, although both dispersal and polyandry might be expected to shape the level of inbreeding depression, this is mainly affected by dispersal, while polyandry has a much more limited effect. These results emphasize the need to consider the potential joint evolution of dispersal and mating system in general, together with their genetic effects, to further our understanding of life-history evolution in spatially structured systems, and provide theoretical expectations for new empirical testing.

## Introduction

Understanding evolution of life-histories and their consequences for populations’ ecology and evolution ultimately requires recognizing the multiple interactions, such as feedbacks, tradeoffs, and shared drivers, existing among different strategies and between potentially competing resolutions of fitness costs. Although most often evolution of different traits is being treated separately, comprehensive understanding of how life-histories evolve under different environmental circumstances, including environmental changes, requires joint evolutionary dynamics to be elucidated. One prominent example is the evolution of competing mechanisms of inbreeding avoidance (Szulkin et al. 2013; Duthie et al. 2018), and specifically the potential for the joint evolution of dispersal and polyandry as competing responses to inbreeding depression, which could then feedback to shape the population’s genetic load and consequent fitness. Yet, such joint evolutionary dynamics have not been examined, precluding comprehensive predictions of mating system evolution in spatially structured populations, as well as its genetic implications.

Inbreeding depression, defined as the reduction in fitness components of offspring of related individuals compared to offspring of unrelated individuals, is a widespread phenomenon that has profound demographic and evolutionary consequences (Keller and Waller 2002; Charlesworth and Willis 2009). It can reduce the mean fitness of a population and increase extinction risk (Theodorou and Couvet 2006; Hedrick and Garcia-Dorado 2016), and it can affect trait evolution (Lande and Schemske 1985; Charlesworth and Charlesworth 1987; Szulkin et al. 2013). Inbreeding depression is widely hypothesized to be a key driver of the evolution of two potential inbreeding avoidance mechanisms, dispersal and polyandry, which play a central role in populations’ ecological and evolutionary dynamics, as they both shape gene flow within and between populations (Waser et al. 1986; Stockley et al. 1993; Perrin and Mazalov 1999; Jennions and Petrie 2000; Tregenza and Wedell 2002). Dispersal, that is any individual movement potentially leading to spatial gene flow (Ronce 2007; Clobert et al. 2012), shapes populations’ spatiotemporal structure as well as their genetic structure, and the extent and direction of gene flow among populations (Clobert et al. 2012). Polyandry, defined as female mating with multiple males within a single reproductive bout (Pizzari and Wedell 2013; Taylor et al. 2014), has only more recently been recognized to have far reaching evolutionary and ecological consequences, and yet remains an evolutionary puzzle (Holman and Kokko 2013; Kvarnemo and Simmons 2013; Pizzari and Wedell 2013). In turn, both dispersal and polyandry can change the relatedness structure within and among populations, thus affecting opportunity for inbreeding and consequent evolution of inbreeding depression (Ronce 2007; Germain et al. 2018).

Despite inbreeding depression being a potential major shared driver, and despite the large amount of both theoretical and empirical work, evolution of dispersal and polyandry given inbreeding have been so far studied separately. Thus, we still do not know whether and how dispersal and polyandry affect each other’s evolution, and how they may feed back onto evolution of inbreeding depression itself. Filling this knowledge gap is particularly important because populations exist in space and it is unlikely that major life-histories, such as dispersal and mating system, evolve independently (Ronce and Clobert 2012; Auld and Rubio de Casas 2013; Hargreaves and Eckert 2014). Further, ongoing environmental changes, such as habitat fragmentation and isolation, are fragmenting populations in smaller demes thus increasing the risk of inbreeding, and more generally demanding understanding of eco-(co-)evolutionary dynamics of life-histories in highly structured systems (Hanski 2011; Cheptou et al. 2017; Legrand et al. 2017).

It is now accepted that inbreeding depression and heterosis (i.e., the increase in fitness in offspring originating from between populations crosses relative to offspring from within population crosses; Charlesworth and Charlesworth 1987; Whitlock et al. 2000; Charlesworth and Willis 2009) can drive dispersal evolution, although debate remains on the form and the relative importance of this effect (Perrin and Goudet 2001; Ronce 2007; Szulkin and Sheldon 2008; Pike et al. 2021). Theoretical work has shown inbreeding depression and heterosis can results in substantial evolution of dispersal, which may be sex-biased or equal between the sexes depending on factors such as the cost of dispersal, the type and strength of same sex competition, the mating system, the strength of inbreeding depression and the presence of demographic and environmental stochasticity (Gandon 1999; Perrin and Mazalov 2000; Guillaume and Perrin 2006, 2009; Roze and Rousset 2009; Henry et al. 2016; Li and Kokko 2019). Substantial insights have been achieved by theoretical models that consider the joint evolution of dispersal and inbreeding depression. These models do not assume constant inbreeding depression but explicitly model the accumulation and purging of deleterious recessive mutations responsible for inbreeding depression and genetic load more generally (Guillaume and Perrin 2006, 2009; Roze and Rousset 2009; Henry et al. 2016). Particularly, Roze and Rousset (2009) by using a continuous chromosome model, which allows modelling a potentially infinite number of deleterious recessive mutations, showed that heterosis can have a much more important effect (relative to kin competition) on dispersal evolution than previously thought (Guillaume and Perrin 2006; Ravigné et al. 2006), especially when population size is large and the genomic deleterious mutation rate is in the upper range of observed values. Further, the effect of heterosis increases when mutations become more recessive (Guillaume and Perrin 2006; Roze and Rousset 2009). However, even studies that include a genetically explicit model of inbreeding depression, generally assume a fixed selection coefficient, *s*, and a dominance coefficient, *h*, across deleterious mutations. Thus, we still do not know how a more realistic distribution of deleterious mutations, which likely comprises many mutations with very small fitness effects and rare ones with larger effects (Eyre-Walker and Keightley 2007), and a negative relationship between selection and dominance coefficients (Agrawal and Whitlock 2011; Huber et al. 2018), might impact on evolving inbreeding depression, and the consequent evolution of dispersal and mating systems (Porcher and Lande 2016).

Meanwhile, explaining the evolution and persistence of polyandry is an ongoing pursuit in evolutionary biology, that is especially challenging when there is direct selection against it, that is when polyandry is costly to females (Arnqvist and Nilsson 2000; Jennions and Petrie 2000; Slatyer et al. 2012; Parker and Birkhead 2013). Although the hypothesis that inbreeding depression can drive the evolution of female multiple mating is prominent among the different evolutionary mechanisms that have been postulated, it remains less established in its theoretical and empirical demonstration, compared to dispersal (Stockley et al. 1993; Jennions and Petrie 2000; Tregenza and Wedell 2002; Reid and Sardell 2012; Reid et al. 2015; Duthie et al. 2016; Germain et al. 2018). Polyandry has been hypothesized to evolve as a mechanism for inbreeding avoidance through two main routes: direct or indirect inbreeding avoidance (Germain et al. 2018). Female multiple mating could evolve because it facilitates inbreeding avoidance through female active pre- and/or post-copulatory allocation of paternity to less closely related males, hence directly reducing inbreeding depression in offspring viability (direct inbreeding avoidance) (Jennions and Petrie 2000; Tregenza and Wedell 2002; Duthie et al. 2016, 2018). Alternatively, without invoking active mate choice and kin recognition, polyandry could evolve because it alters relatedness among the female’s offspring, producing more half-sibs rather than full-sibs, thereby reducing the risk of close inbreeding for the offspring of a polyandrous female and reducing inbreeding depression in her grand offspring (indirect inbreeding avoidance) (Cornell and Tregenza 2007; Germain et al. 2018). The few theoretical models to have investigated these verbal predictions (Cornell and Tregenza 2007; Duthie et al. 2016, 2018) have generally concluded that the strength of indirect selection on polyandry through inbreeding avoidance might be very small, thereby suggesting a minor role of inbreeding depression in polyandry evolution.

Specifically, Cornell and Tregenza (Cornell and Tregenza 2007) concluded that purging of deleterious recessive mutations makes it unlikely to maintain sufficient levels of inbreeding depression to favor costly polyandry, and that the evolution of polyandry as a mechanism of indirect inbreeding avoidance is far more likely if inbreeding depression was due to overdominance. This poses a problem because current understanding suggests that inbreeding depression is predominantly caused by deleterious recessive, rather than overdominant, mutations (Charlesworth and Willis 2009). However, Cornell and Tregenza’s (2007) model makes some assumptions that preclude assessing whether such a mechanism could generally drive polyandry evolution in spatially structured populations. The model considers alternating generations of outbreeding and inbreeding, which is particularly relevant to some invertebrate groups where mated females cyclically colonize empty patches, thus experiencing cyclical changes in inbreeding risk (e.g., store product pest species such as flour beetles, *Tribolium* spp.). However, this does not apply to species with more regular inbreeding as it does not consider the building up of complex relatedness structure within a population, that arise across multiple generations and might weaken the selective advantage of polyandry (Germain et al. 2018). Indeed, Cornell and Tregenza’s (Cornell and Tregenza 2007) pointed out that higher levels of inbreeding, as we might expect in spatially structured populations, might favor polyandry, although this might be offset by greater purging of deleterious recessives alleles. No model so far has investigated the evolution of polyandry as a mechanism of indirect inbreeding avoidance in populations that are spatially structured and connected by dispersal, nor has considered an explicit model of inbreeding depression with realistic distributions of the fitness effects and dominance coefficients of deleterious mutations, thus leaving a substantial knowledge gap.

Beyond the unknowns that are still present in the separate theories of dispersal and polyandry evolution given inbreeding depression, we do not know how these two potential mechanisms of inbreeding avoidance might affect each other’s evolution, and feed back onto evolution of inbreeding depression. I hypothesize a negative feedback between dispersal and polyandry, whereby the evolution of polyandry might reduce inbreeding and hence reduce the strength of selection for dispersal and, *vice versa*, the presence of high dispersal might weaken selection for polyandry. The outcome of this tug-o-war will likely depend on the relative efficiency of dispersal and polyandry in reducing inbreeding, with the expectation that dispersal would be much more effective, on the level of inbreeding load, and on the strength of direct selection against them, that is on the cost of dispersal and polyandry.

I investigate this hypothesis with a modelling framework that allows joint evolution of dispersal and polyandry in spatially structured populations and, at the same time, explicit accumulation of deleterious mutations and evolution of inbreeding depression. Specifically, inbreeding depression is determined by accumulation of a potentially infinite number of deleterious recessive mutations with a realistic distribution of fitness effects and dominance coefficients. First, I test whether existing predictions on the independent evolution of dispersal and polyandry hold in spatially structured populations given an explicit model of inbreeding and inbreeding depression evolution, and how their evolution affects inbreeding depression. Second, I test the novel hypothesis of a negative feedback between jointly evolving dispersal and polyandry and, third, determine their joint effect on the evolution of inbreeding depression. More generally, I show the value of moving towards theoretical frameworks that explicitly integrate ecology, genetics, and evolution, to progress our understanding of life-history evolution and its impacts on populations.

### The Model

To investigate the joint evolution of dispersal and polyandry given inbreeding depression I built a spatially and genetically explicit individual-based model where emigration probability (*d*) and female re-mating rate (*a*) evolve. Genetic load and resulting inbreeding depression (ID) also evolve by accumulation and purging of deleterious recessive mutations. Populations of a dioecious species, with non-overlapping generations, occupy cells within a landscape grid of 20 by 20 cells, and are connected by dispersal. The environment is spatially homogeneous and temporally constant; each cell is suitable to hold a population with constant carrying capacity *K =* 50. All the model variables and parameters are listed in Table S1.

### Genetic architecture and inbreeding depression

To model the genetic basis of *d* and *a*, individuals carry two unlinked diploid loci with a continuous distribution of alleles (Kimura 1965). The initial value of each allele is sampled from normal distributions. Alleles can mutate with probability *μ* = 10^-3^/allele/generation. When a mutation occurs a random normal deviate with mean zero is added to the allele value. The individual’s genotypic values for the two traits, *g_d_* and *g_a_*, are given by the sum of the two allelic values at the respective loci. The phenotypic expression has no environmental variance and is female limited for *a*. I assume the phenotypes *a* ≥ 0 and 0 ≤ *d* ≤ 1.

ID is determined by deleterious recessive mutations (Charlesworth and Willis 2009) which accumulate on a continuous chromosome (Roze and Rousset 2009). Each individual carries two homologous autosomes of length *R* (genome map length). The position of each new deleterious mutation on the chromosome is sampled from the continuous uniform distribution *U*[0, *R*]. The number of loci at which mutations can occur is therefore effectively infinite (“infinite site model” (Peischl et al. 2015)). Each new mutation is characterized by a selection coefficient *s*, determining the mutation’s effect in the homozygous state, and a dominance coefficient *h*. The effect of each mutation *i* is multiplicative such that the genetic fitness *ω* of an individual is given by

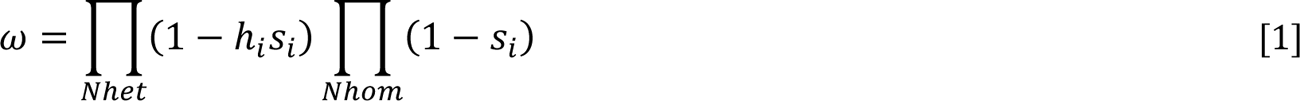

*Nhet* and *Nhom* represent the number of heterozygous and homozygous mutations respectively. Deleterious mutations are of two types: mildly deleterious and lethal (Gilbert et al. 2017; Spigler et al. 2017). Mildly deleterious mutations occur at a rate *U_d_* = 1.0/diploid genome/generation (Haag-Liautard et al. 2007; Zhu et al. 2014). The selection coefficient of each new mutation is sampled from a gamma distribution with mean *s_d_* = 0.05 and shape parameter *α* = 1 (Schultz and Lynch 1997; Spigler et al. 2017). The dominance coefficient of a mutation *i* depends on its selection coefficient *s_i_* and is sampled from the continuous uniform distribution *U*[0.0, e^−KS_i_^]. *k* is defined as − log(2ℎ_d_)⁄S_d_, where *h_d_* is the mean dominance coefficient (*h_d_* = 0.3) (Caballero and Keightley 1994; Spigler et al. 2017). Lethal mutations occur at rate *U_l_* = 0.2/diploid genome/generation and are extremely recessive, with constant selection coefficient *s_l_* = 1 and dominance coefficient *h_l_* = 0.02 (Simmons and Crow 1977; Lande et al. 1994; Porcher and Lande 2005; Spigler et al. 2017). At each generation, the number of new mutations per diploid genome is sampled from Poisson distributions with parameters *U_d_* and *U_l_*. The number of crossovers along the continuous chromosomes is sampled from a Poisson distribution with mean *R*, and the position of each crossover is sampled from the uniform distribution *U*[0, *R*].

Individuals also carry *L_n_* = 500 neutral autosomal diploid loci to determine the degree to which individuals are inbred (Bocedi and Reid 2017). Neutral allelic values are continuously distributed, sampled from the uniform distribution *U*[−1000.0, 1000.0], and mutate with probability 10^-3^ /allele/generation. Neutral loci recombine at rate *r* = 0.1. Alleles at the same locus will be identical only by descent as the chance of non-descent identity by state, stemming from initialization or mutation, is negligible. For this reason, individual’s neutral homozygosity, defined as the number of neutral homozygous loci / L_n_, represents a proxy for the realized individual coefficient of inbreeding, hereafter noted as *F_homoz_* (Markert et al. 2004; Neff and Pitcher 2008; Fromhage et al. 2009; Bocedi and Reid 2017). When an individual is born, it is assumed to die immediately if its genetic fitness *ω* = 0. If *ω* > 0, the newborn survives to adulthood (unless it incurs in dispersal mortality, see below) when *ω* will determine its probability of reproducing. Thus, ID is affecting two fitness components: 1) reduction in offspring survival, which is determined by the presence of homozygous lethal mutations, and 2) reduction in adult probability of reproducing, which is determined mainly by mildly deleterious mutations.

The level of ID present in a metapopulation at a given point in time, was calculated as: 1) effect of *F_homoz_* on offspring survival by fitting a generalized linear model with Poisson distribution and logarithmic link function (Nietlisbach et al. 2019); 2) effect of *F_homoz_* on the logarithm of adult reproduction probability by fitting a linear model (Morton et al. 1956). All models were fitted in R (R Core Team 2019). To obtain a spread of *F_homoz_* values to estimate ID, I created individuals by selecting the central 140 populations in the landscape (out of 400 populations) at a given point in time. For each of these populations, I mated each female with 10 males selected randomly within the female’s population and with 10 males selected randomly between different populations. Each mating produced one offspring. Models were then fitted on all the offspring pooled together. These individuals were also used to calculate the level of heterosis (H) in each population as H = 1 − ω_w_⁄ω_b_, where ω_w_ is the genetic fitness of offspring produced within populations, and ω_b_ is the genetic fitness of offspring produced between populations (e.g., Roze and Rousset 2009). All the offspring produced this way were used only for estimating ID and H, and then discarded, thus were not part of the ecological and evolutionary dynamics.

### Life cycle and selection

At each generation, the life cycle consists of reproduction (mating and offspring birth), adults’ death and death of offspring with *ω* = 0, offspring dispersal and density-dependent survival. An adult’s probability of reproducing is given by its genetic fitness *ω*. Each reproducing female *i* mates initially once, and then re-mates with probability *Pmat_i_* depending on her re-mating rate phenotype *a_i_* and on her current number of mates *Nm_i_*:

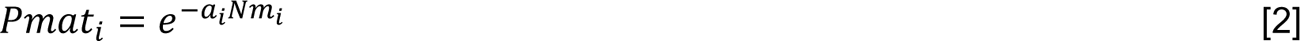

Each mate is randomly sampled among the reproducing males in the female’s population, without replacement. If the female has already mated once with all the reproducing males in the population, she stops re-mating. Mating multiply can be costly to females. The probability that a female *i* survives to reproduction (*ψ_i_*) depends on her total number of mates *Nmates_i_* and on the strength of selection against multiple mating ω^2^:

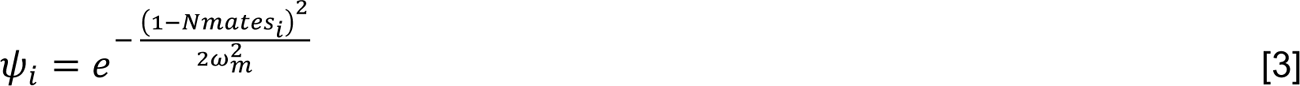

If the female survives mating, she produces a number of offspring sampled from a Poisson distribution with mean *f* = 12 and primary sex-ratio = 1:1. Each offspring is sired by a male randomly chosen, with replacement, between the female’s mates.

After reproduction, all adults die, and offspring may disperse among sub-populations according to their emigration probability phenotype *d*. Dispersal distance and direction are sampled from a negative exponential distribution (mean 2 cells), and uniform distribution between 0 and 2π, respectively. The new location is re-sampled if it falls outside the grid. Dispersal has a cost, *c_d_*, representing the probability of an individual dying during dispersal. After dispersal, density-dependent survival takes place in each population. Individuals survive with probability *min*(K⁄N, 1), where *N* is the total number of individuals in the population.

### Simulations

I ran three main sets of simulations. 1) Only dispersal is evolving while female re-mating rate is constant, *a* = 3 (corresponding to 1.3 mates per female on average – hereafter defined as monandry); 2) only female re-mating rate is evolving while dispersal is constant, *d* = 0.05; 3) both dispersal and female re-mating rate are evolving. All simulations were run under varying costs of dispersal and female re-mating and repeated in the absence of deleterious mutations (and hence ID) as control. I additionally tested the effect of varying the rates of deleterious mutation (*U_d_* = 0.5 and *U_l_* = 0.1; *U_d_* = 0.1 and *U_l_* = 0.02).

## Results and Discussion

### Evolution of dispersal and inbreeding depression given monandry

Under fixed monandry conditions, emigration probability *d* evolved in response to the presence of deleterious recessive mutations and consequent inbreeding depression (ID), reaching higher values the lower the cost of dispersal (*c_d_*; Fig. 1A). Much higher emigration probability evolved in the presence than in the absence of genetic load when, given the spatio-temporally homogeneous environment, the only driver of dispersal evolution was kin competition. For example, at the lower cost considered (*c_d_* = 0.3), *d* evolved almost eight times higher with ID [median d̅ = 0.41 (95%CI 0.401-0.414), where d̅ represents the mean phenotypic value for one replicate metapopulation; median and CI are taken across 20 replicates] than without ID [median d̅ = 0.05 (95%CI 0.05-0.054)]; while it was almost seven times higher at high cost of dispersal [*c_d_* = 0.6; median d̅ = 0.15 (95%CI 0.15-0.157) with ID *vs*. median d̅ = 0.02 (95%CI 0.02-0.023) without ID]. Results remained qualitatively similar under lower rates of deleterious mutations, although the lower the mutation rate the lower the evolved dispersal probability (Fig. S1A-S2A). The level of heterosis H reached at equilibrium depended on the level of evolved dispersal, and hence on dispersal cost (Fig. S3A). Higher cost of dispersal led to lower dispersal and to populations being less homogeneous in terms of their genetic load, and hence to higher heterosis. Higher rates of deleterious mutations, by causing accumulation of higher genetic load, led to the emergence of higher heterosis compared to lower mutation rates. These results broadly agree with the conclusions of Roze and Rousset (2009) who, by using a genetically explicit, infinite site model of genetic load, showed that heterosis can have a substantial role in dispersal evolution, in contrast with what shown by previous models (Guillaume and Perrin 2006; Ravigné et al. 2006). Importantly, the results also show that the qualitative predictions made by (Roze and Rousset 2009) hold when assuming a realistic distribution of mutational effects in terms of selection and dominance coefficients; despite the accumulation of many mutations with very low (almost neutral) selection coefficient, substantial heterosis is maintained as it is its substantial role (relative to kin competition) in driving dispersal evolution.

**Figure 1.**
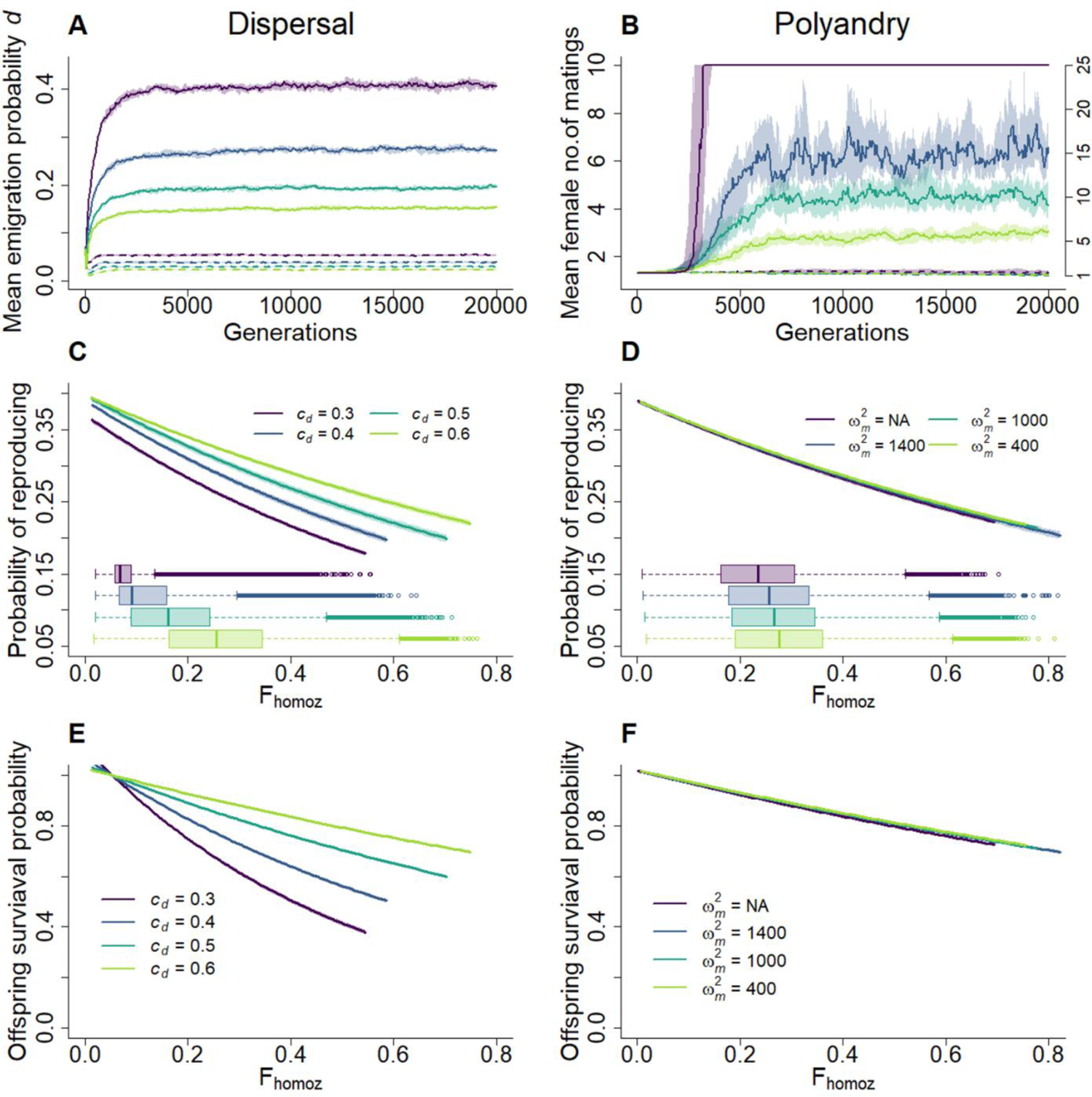
Inbreeding depression promotes evolution of dispersal and polyandry when either one trait or the other evolves. **A)** Evolution of mean dispersal probability phenotypes *d* in the absence of polyandry (*a* = 3.0), under different costs of dispersal (*c_d_* = 0.3, 0.4, 0.5, 0.6), in the absence (dashed lines) or presence (solid lines) of deleterious mutations. **B)** Evolution of mean polyandry phenotypes (expected female number of matings *P* = 1 + 1/*a*) under fix dispersal probability (*d* = 0.05), as a function of different strengths of direct selection against female remating (no cost; ω^2^= 1400, 1000, 400) in the absence (dashed lines) or presence (solid lines) of deleterious mutations. In the presence of deleterious mutations and no direct selection against polyandry (ω^2^ = NA), females evolve to mate with all the males in the population; the y-axis on the right hand-side refers to this single line (purple). Lines represent the median of mean phenotypes across 20 replicated simulations; colored shades depict the first and third quartile. **C-D)** Relationship between individual probability of reproducing and inbreeding coefficient *F_homoz_* (i.e., ID in reproduction probability) when C) dispersal evolves under different costs, in the absence of polyandry and, D) polyandry evolves under different strengths of direct selection, with fix dispersal probability. Lines show the fitted models and colored shades the 95% CI. Models are fitted at generation 20,000 to a subsample of 140 populations, across 10 replicates. Boxplots represents the distribution of the individual *F_homoz_*. **E-F)** Relationship between offspring survival probability and *F_homoz_* (i.e., ID in offspring survival probability). In E) simulation scenarios and parameters as in C); in F) as in D). The coefficients (i.e., mutation load and inbreeding load) and standard errors of all the fitted models are presented in Table S2.

The level of evolved ID depended on the level of evolved dispersal, and hence on dispersal cost (Fig. 1C-E; Table S2). Higher dispersal cost led to lower *d* and consequent higher inbreeding within local populations, reflected by higher neutral homozygosity (Fig. 1C). In turn, more inbreeding facilitated purging of deleterious recessive mutations thus reducing ID. This was true for both components of ID. Both lethal mutations (strongly recessive mutations causing ID in early life – offspring survival, Fig. 1E) and mildly deleterious mutations (causing ID in later life – adult reproduction probability, Fig. 1C) experienced greater purging at lower dispersal, where both the mutation load (i.e., the decrease in fitness for outbred individuals) and ID were lower compared to scenarios with much higher dispersal evolving, albeit grater differences were observed for ID rather than for mutation load (Table S2). This pattern was conserved at lower mutation rates although much less genetic load accumulated, leading to higher individual fitness and lower mutation and inbreeding load (Fig. S1C-E; S2C-E). There is therefore a positive feedback between evolution of dispersal and inbreeding depression, whereby high dispersal maintains high levels of inbreeding depression, which in turn maintains positive selection for dispersal.

### Evolution of polyandry and inbreeding depression given low dispersal

Like dispersal, polyandry (expected female number of matings, *P* = 1 + 1/*a*) evolved in response to ID when dispersal was low and not evolving, to a level that depended on the fitness cost of female multiple mating (Fig. 1B; Fig. S4). Given no cost of polyandry, females evolved to mate with all the males present in the population. However, even a very small cost reduced substantially P̅ (where P̅ represents the mean phenotypic value for one replicate metapopulation); yet moderate polyandry evolved. For example, strength of direct selection on re-mating ω^2^= 1000 led to evolution of median P̅ = 4.13 (95%CI 3.51-5.26) (median and CI are taken across 20 replicates), corresponding to an average realized survival cost of 0.005 for females. On the contrary, in the absence of ID, polyandry did not evolve even when free of cost (Fig. 1B and S4, dashed lines). Lower rates of mildly deleterious and lethal mutations substantially reduced selection for polyandry such that hardly any polyandry evolved, or evolution took considerably longer time (Fig. S1B; S2B).

These results provide a positive answer to the still standing question of whether costly polyandry can evolve, at least in theory, as a mechanism of indirect inbreeding avoidance in spatially structured populations (Germain et al. 2018). Specifically, they show that costly polyandry, although the cost needs to be quite low, as predicted by (Cornell and Tregenza 2007), can indeed evolve in response to inbreeding depression given: i) inbreeding depression caused exclusively by deleterious recessive mutations, and without the need for overdominant mutations (Cornell and Tregenza 2007); ii) realistic deleterious mutation rates and distribution of fitness effects (Caballero and Keightley 1994; Schultz and Lynch 1997; Haag-Liautard et al. 2007); iii) and complex sibship structure emerging in spatially structured populations (Germain et al. 2018). Thus, polyandry evolution as a means of indirect inbreeding avoidance might be more widely spread than previously thought (Cornell and Tregenza 2007; Germain et al. 2018), and therefore a potentially important mechanism to explain the existence of even low levels of polyandry across multiple systems, especially when dispersal is low.

Although polyandry evolved in response to the presence of genetic load, the level of evolved polyandry under different costs did not substantially change the level of inbreeding (measured as neutral homozygosity) and ID at equilibrium (Fig. 1D-F; Table S2; Fig. S1D-F; S2D-F). In fact, both the mutation load and ID were the same across levels of evolved polyandry and polyandry’s costs. The level of mutation and inbreeding load was mainly determined by dispersal, which was fixed to *d* = 0.05, and similar to what evolved under high cost of dispersal and no polyandry (Fig. 1C-E). This was true also for heterosis (Fig. S4). Thus, under low fixed dispersal probability, polyandry seems to have a minimal effect, if any, on the purging or accumulation of genetic load, ID and, not surprisingly, heterosis. Unlike with dispersal therefore, with polyandry there seems not to be scope for a positive feedback with inbreeding depression, but just for one-direction effect of inbreeding depression on polyandry evolution.

### Joint evolution of dispersal, polyandry, and inbreeding depression

When dispersal and polyandry could evolve jointly, results clearly confirmed the hypothesis of a negative feedback between two competing mechanisms of inbreeding avoidance (Fig. 2; S5). This feedback was modulated by the relative costs of the two evolving behaviors. For a given cost of dispersal (*c_d_*), higher dispersal evolved under monandry; *vice versa*, the higher the evolved level of polyandry (hence for lower costs of female re-mating) the lower the evolved emigration probability. For example, *c_d_* = 0.3 led to evolution of median d̅ = 0.39 (95%CI 0.375-0.396) and median P̅ = 1.97 (95%CI 1.763-2.173) for high cost of female re-mating (ω^2^ = 400), while it led to median d̅ = 0.31 (95%CI 0.312-0.322) and to females mating with all the males in the population (*P* = 25) when female re-mating did not carry costs. On the other hand, for a given cost of female re-mating, higher polyandry evolved given lower evolved dispersal (hence for higher *c_d_*). For example, ω^2^= 1000 led to evolution of median d̅ = 0.13 (95%CI 0.126-0.131) and median *P̅* = 4.49 (95%CI 3.842-5.317) for high cost of dispersal (*c_d_* = 0.6), while it led to median d̅ = 0.38 (95%CI 0.36-0.383) and median *P̅* = 2.58 (95%CI 2.467-2.814) for *c_d_* = 0.3. This joint evolution and negative feedback did not emerge when the two traits evolved in the absence of genetic load and hence ID (Fig. S6), where dispersal evolved to very low levels while polyandry did not evolve.

**Figure 2.**
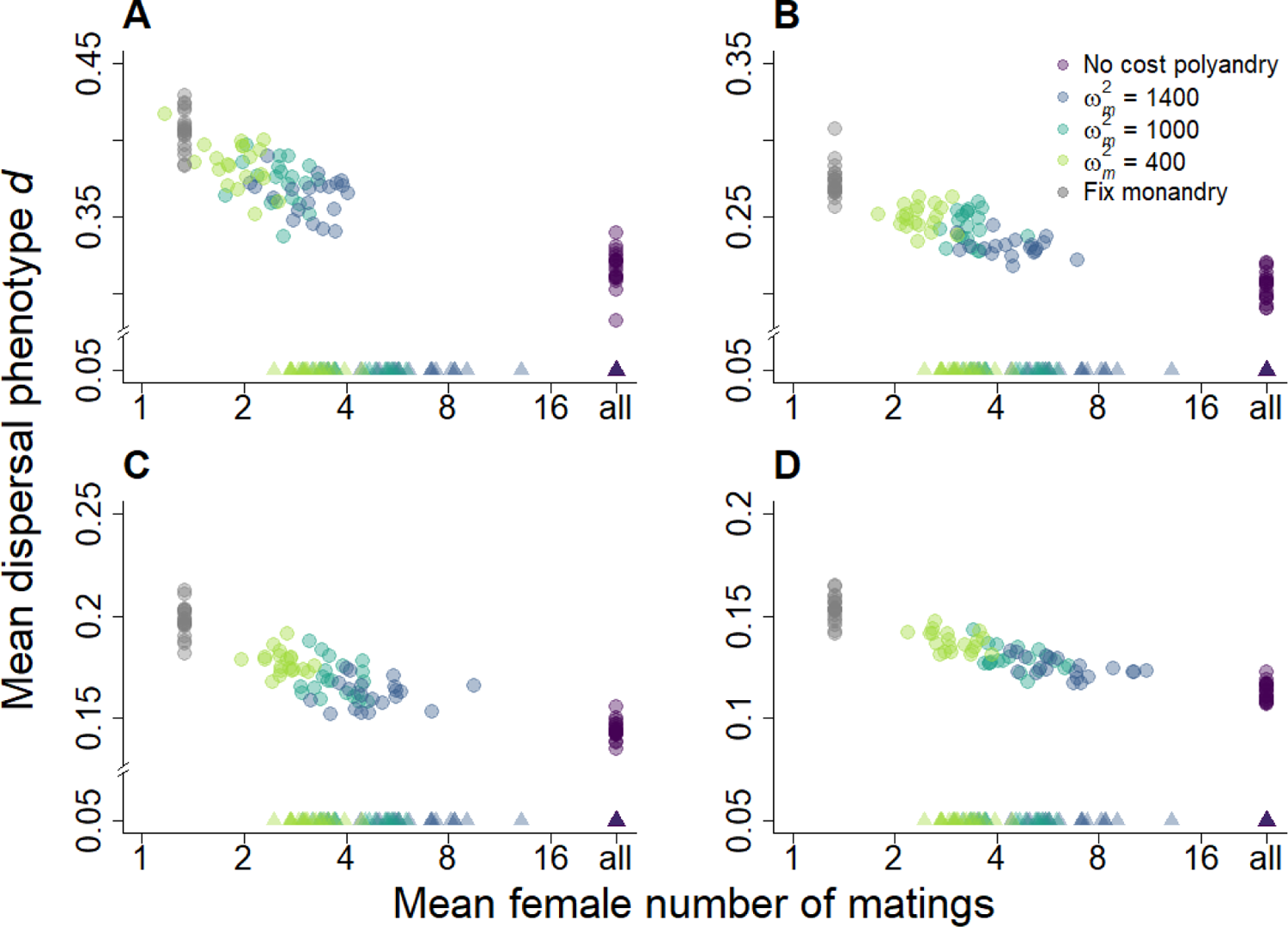
Dispersal and polyandry negatively feedback to each other evolution under inbreeding depression. Joint evolution of mean dispersal probability phenotypes *d* and mean polyandry phenotypes (mean female number of matings: *P* = 1 + 1/*a*) in the presence of inbreeding depression, given different costs of dispersal *c_d_* (**A:** 0.3; **B:** 0.4; **C:** 0.5; **D:** 0.6) and different strengths of direct selection against female re-mating (no cost; ω^2m^ = 1400, 1000, 400). Each data point represents the mean phenotypic values for one out of 20 replicate simulations at generation 20,000. Colored dots indicate simulations where dispersal and polyandry jointly evolved; triangles, simulations where polyandry evolved given fix dispersal probability (*d* = 0.05); grey dots, simulations where dispersal evolved given fix monandry (*a* = 3.0). The x-axis is on the logarithmic scale to aid visualization. Note the different y-axis scales for the four panels. Other parameters: *U_d_* = 1.0, *U_l_* = 0.2.

The negative correlation between evolved emigration probability (*d*) and polyandry (*P*) was present at the metapopulation level (i.e., simulation replicate level), whereby for a given cost of dispersal and strength of direct selection on female re-mating, systems that evolved higher emigration probability phenotypes, also evolved lower polyandry, and *vice versa* (Fig. S7). This correlation was present when female re-mating was costly; in the absence of re-mating costs the correlation disappeared as females consistently evolved to mate with all the males in the population. Within a metapopulation, there was no evidence of a negative correlation between *d* and *P*, whereby subpopulations with higher dispersal might have had lower polyandry (Fig. S8). Further, there was no evidence of any genetic correlation between *g_d_* and *g_a_*, nor between *d* and *P* (Fig. S9).

Lower rates of mildly deleterious and lethal mutations led to evolution of lower dispersal and polyandry (Fig. S10), similarly to when the two traits evolved independently (Fig. S1-S2). Polyandry hardly evolved, apart from a few exception simulations under no or weak direct selection against female re-mating, where it reached high levels. A strong negative correlation between dispersal and polyandry phenotypes was present at the metapopulation level when some polyandry evolved, while it disappeared for very low mutation rates combined with costly polyandry (Fig. S11). As for higher rates of deleterious mutations, at lower mutation rates there was no evidence of a negative correlation between dispersal and polyandry at the subpopulation level, nor of any genetic correlation (Fig. S12-S13).

The evolved level of ID depended mainly on the cost of dispersal, and hence on the level of evolved dispersal (Fig. 3-4; Table S3). As for the scenario where dispersal evolved under fixed monandry (Fig. 1C-E), higher cost and consequent lower evolved dispersal led to lower inbreeding depression to both probability of reproducing (Fig. 3A, Fig. 4 diamonds) and offspring survival (Fig. 3B, Fig. 4 dots). Interestingly, under the joint evolution of dispersal and polyandry, polyandry had a slight but detectable effect on the level of evolved ID, compared to when evolving under fixed low dispersal (Fig. 1D-F). Specifically, for a given dispersal cost, higher polyandry (lower strength of direct selection on female re-mating) led to the accumulation of lower inbreeding load in offspring survival, especially for lower costs of dispersal (Fig. 4 dots; Table S3). The difference was especially evident when comparing no cost *vs* costly polyandry. For the inbreeding load in adult reproduction, this effect was present but only very slight and with overlapping confidence intervals between different costs of polyandry (Fig. 4 diamonds; Table S3). This result is perhaps surprising and counterintuitive as the expectation would be for more polyandry to lead to accumulation of higher load, due to reduced inbreeding and purging, and not the opposite as observed here. However, this effect might be due to high polyandry reducing evolved dispersal thus, in fact, facilitating slightly greater purging through its effect on dispersal.

**Figure 3.**
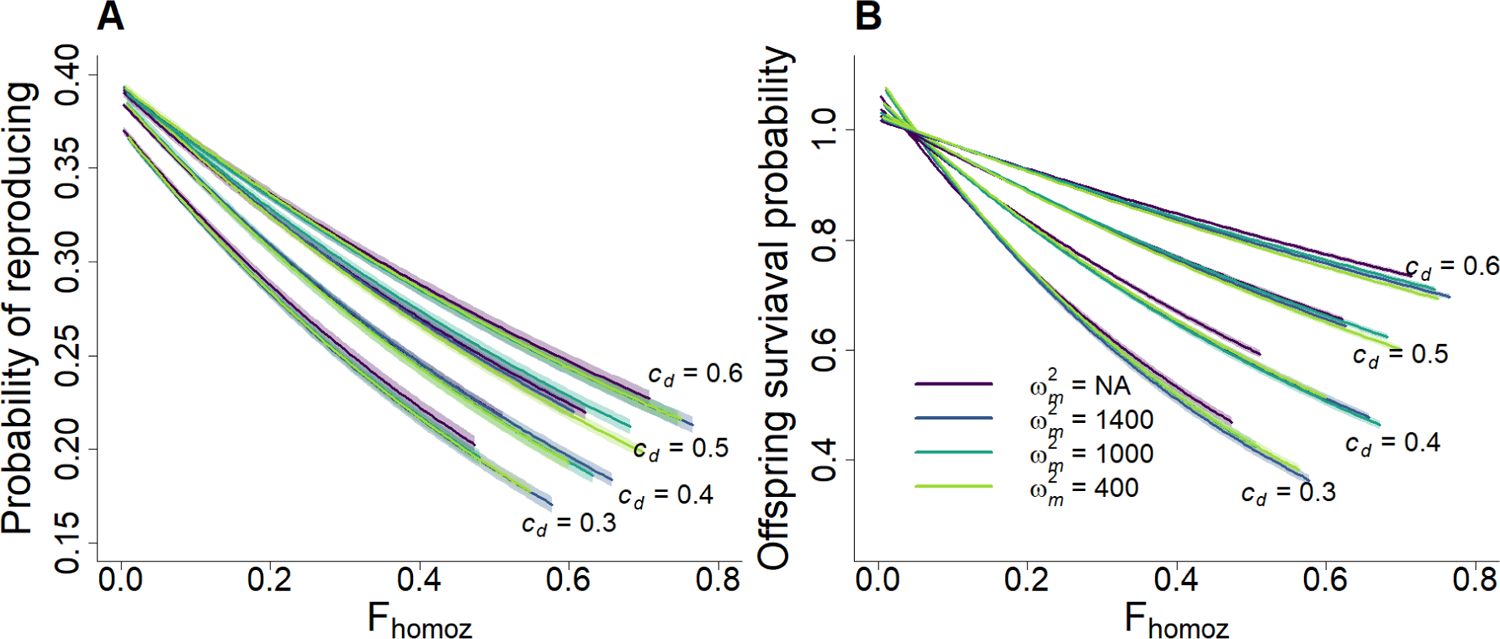
Jointly evolving dispersal and polyandry affect evolution of inbreeding depression in reproduction and survival. Relationship between **A)** individual probability of reproducing and neutral homozygosity F_homoz_ (i.e., inbreeding depression in reproduction probability), and **B)** between offspring survival probability and neutral homozygosity (i.e., inbreeding depression in offspring survival probability). Results are presented for varying costs of dispersal (*c_d_*) and strengths of direct selection against female multiple mating (ω^2^_m_; colors). Lines show the fitted models and colored shades the 95% CI. The coefficients (i.e., mutation load and inbreeding load) and standard errors of all the fitted models are presented in Table S3. Models are fitted at generation 20,000 to a subsample of 140 populations, across 10 replicates.

**Figure 4.**
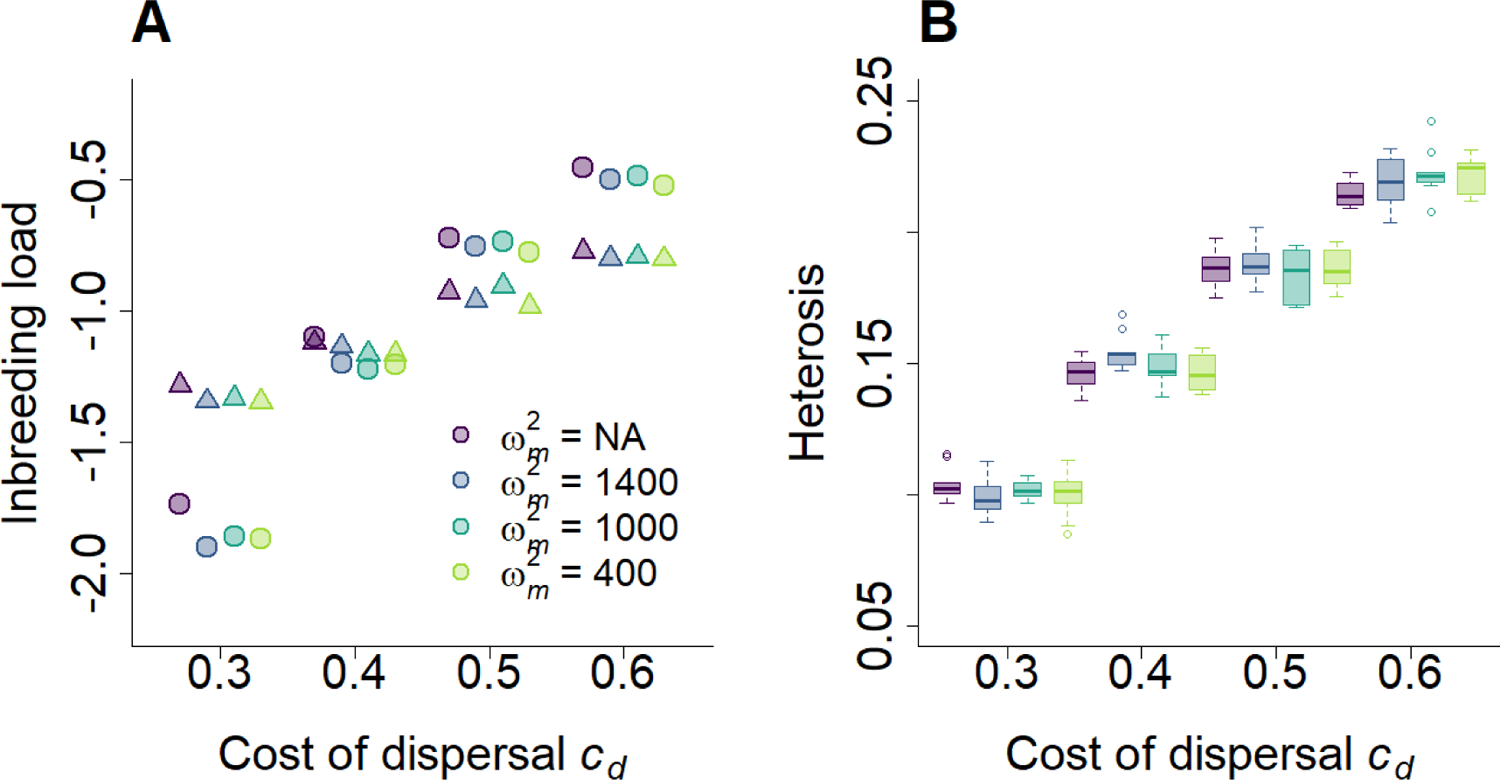
Inbreeding load and heterosis emerging under the joint evolution of dispersal and polyandry. **A)** Estimated slopes of probability of offspring survival (dots) and probability of reproduction (triangles) on individual neutral homozygosity F_homoz_ (i.e., inbreeding load), for different cost of dispersal (*c_d_*) and strengths of direct selection on female multiple mating (ω^2^_m_; colors). Results are presented on the log scale. Standard errors are not shown because smaller than the dots size. Models are fitted at generation 20,000 to a subsample of 140 populations, across 10 replicates. **B)** Heterosis as a function of *c_d_* and ω^2^_m_ (the color legend is the same as in A). Heterosis is shown as median (solid bands), first and third quartiles (box limits), and approximately twice the standard deviation (whiskers) over 20 replicate simulations at generation 20,000.

Finally, the emerging heterosis in the system was exclusively driven by the level of evolved dispersal, and hence by dispersal cost. Higher heterosis was present under high costs of dispersal, thus generating strong positive selection for dispersal, counteracting the high cost (Fig. 4B).

### General discussion

These results shed light on the previously unconsidered intimate connection existing between dispersal and polyandry evolution through their shared driver of inbreeding depression, and on their effect on evolution of inbreeding depression itself. They highlight important interactions between the evolution of two fundamental life-histories which shape gene flow in space and time, and thereby affect species’ eco-evolutionary dynamics. More broadly, these results demonstrate the need to consider life-history evolution as happening within a complex and integrated system, where multiple competing and degenerate routes to reduce fitness costs can arise and affect each other’s evolutionary dynamics (Edelman and Gally 2001; Mason 2015). This becoming strongly apparent in different complex biological systems, such as genetic codes and networks, neural networks, organismal development, population and community dynamics (see examples within Edelman and Gally 2001; Mason 2015)), but perhaps it has not been into the investigation of life-history evolution. For example, within the context of evolution of polyandry as inbreeding avoidance strategy, Duthie et al. (2018) showed that evolution of pre-copulatory and post-copulatory mechanisms of inbreeding avoidance and associated polyandry is affected by evolutionary feedbacks and degeneracy. Thus, understanding when and how we can expect to observe different patterns of life-history traits co-occurrence requires explicitly modelling feedbacks between competing and / or complementary evolutionary routes.

Here, I show that through their effect on the population relatedness structure, and hence on inbreeding levels, dispersal and polyandry can negatively feedback to each other evolution, thereby providing two competing mechanisms for compensating the negative fitness effects of genetic load. The engine of this feedback is inbreeding depression. Without genetic load giving rise to inbreeding depression and heterosis, kin competition alone is not sufficient for a negative relationship between dispersal and polyandry to emerge, nor for polyandry to evolve. Despite the recognition that dispersal and mating system evolution may be profoundly interconnected, leading to observable dispersal-mating system syndromes (Ronce and Clobert 2012; Auld and Rubio de Casas 2013; Hargreaves and Eckert 2014), previous theory has rarely focused on their joint evolution, with the notable exception of joint evolution of dispersal and self-fertilization (Massol and Cheptou 2011; Sun and Cheptou 2012; Iritani and Cheptou 2017). Further, the relatively recent recognition of the widespread occurrence and importance of female multiple mating meant that the relationship of this key component of many mating systems with dispersal has been understudied compared to monogamy and polygyny. Probably the paucity of theory and clear testable predictions, and the inherent difficulties in studying these two complex suits of behaviors empirically (Rhainds 2017), underlie the scarcity of empirical evidence on patterns of dispersal-polyandry co-occurrence and joint evolution, especially, but not exclusively, at the individual level (Laloi et al. 2009; Reid and Arcese 2020).

This model provides the clear prediction of a negative correlation between dispersal and female multiple mating at the species or metapopulation level, provided the presence of inbreeding depression. This means that we might expect to find that females of highly dispersive species (or metapopulations) engage less in multiple mating, while expecting to observe high frequency of female multiple mating in highly philopatric species (or metapopulations) for which, for example, dispersal is very costly. A recent analysis of song sparrow’s (*Melospiza melodia*) long term pedigree data from the population occupying the small island of Mandarte (British Columbia, Canada) showed that recent immigrants to the population had lower breeding values for female extra-pair reproduction (which results from underlying polyandry) than the local population (Reid and Arcese 2020). This rare empirical evidence points towards a negative correlation between dispersal and female multiple mating, in this case present between the island and the mainland (much larger) population, although the causes of such relationship are currently unknown.

Results from my current model did not show any covariance between polyandry and dispersal at finer spatial scale (e.g., within a metapopulation), or at the individual level. Likewise, very few empirical studies have explicitly tested for the presence of such covariance, and not always found evidence of it (Rhainds 2017; Rafter et al. 2018; Reid and Arcese 2020). However, there are assumptions embedded within the model structure that could affect these predictions, pointing towards the need for further theoretical work needed to resolve whether we should expect any covariance between these two traits at the population or individual level. The model is ecologically quite simple; the environment is spatially homogeneous as are the costs of dispersal and polyandry. Similarly, the relatively high levels of dispersal evolving, combined with the long-distance dispersal allowed by the negative exponential dispersal kernel and the homogeneous environment (and consequently homogeneous risk of inbreeding), impede populations to diverge in their evolutionary trajectories for dispersal and polyandry, such that populations evolve as a single system despite the emerging internal relatedness structure. A spatially heterogeneous environment where, for example, some populations are more isolated than others, or where populations experience different costs of dispersal and/or polyandry, may promote a negative relationship between dispersal and polyandry at the population level whereby frequency of female multiple mating may be expected to be higher in more isolated populations.

The model does not allow for evolution of sex-biased dispersal (Li and Kokko 2019). Although evolution of sex-biased dispersal is not a necessary condition for inbreeding avoidance through dispersal (Guillaume and Perrin 2009; Roze and Rousset 2009; Li and Kokko 2019)and, on the other hand, it is itself affected by multiple ecological and evolutionary drivers other than inbreeding depression (Henry et al. 2016; Li and Kokko 2019), evolution of sex-biased dispersal could change the dynamics of joint evolution of dispersal and polyandry in ways that are difficult to predict without explicit investigation. Evolution of sex-biased dispersal and the direction of the bias depend on sex differences in fitness variance between patches, where, generally, the sex with the larger between-patch variance in fitness evolves to disperse more (Li and Kokko 2019). As the effect of polyandry on sex-specific variance in fitness is hard to predict (Lotterhos 2011; Bocedi and Reid 2017), this cascades in the challenge to predict its effect on sex-biased dispersal. Interestingly, although previous models considered the effect of different mating systems on the evolution of sex-biased dispersal given inbreeding depression (Hirota 2005; Guillaume and Perrin 2009; Henry et al. 2016), no model to my knowledge has considered the opposite, that is the effect of sex-biased dispersal on mating system evolution, nor the potential feedbacks between the two, revealing yet another knowledge gap in our understanding of life-history evolution in spatially structured systems.

The life cycle represented in this model is also relatively simplified with mating occurring after dispersal and non-overlapping generations. The timing of mating relative to dispersal can affect gene flow and the genetic structure of the population thereby affecting evolution of both dispersal and mating system (Hirota 2004; Shaw and Kokko 2015; Lakovic et al. 2017). In species with post-mating dispersal, as many invertebrate species where females disperse after mating and before oviposition, polyandry has been hypothesized to be particularly beneficial for dispersing females colonizing new habitat patches as their offspring would benefit from half-sib rather than full-sib matings, thus reducing the level of inbreeding in the females’ grand-offspring (Cornell and Tregenza 2007). Moreover, post-mating dispersal is much less effective, if at all, in avoiding inbreeding as mating takes place within the natal population (Li and Kokko 2019). In this case, a positive covariance between dispersal and polyandry would be expected. Although high levels of polyandry have been demonstrated in the field for two stored grain pest beetles with post-mating dispersal, a positive covariance between dispersal and female multiple mating has not been found (Rafter et al. 2018). This positive association between polyandry and dispersal in species with post-mating dispersal could be particularly beneficial during colonization of empty patches thus, for example during range expansion or shifting, relative to within an established spatially structured population (Rafajlović et al. 2013; Ding et al. 2017). Indeed, experimental studies with seed and flour beetles have shown that multiple mated females establish fitter populations both in benign and thermally challenging environments (Power and Holman 2014; Lewis et al. 2020). These results, together with the ones presented here, point to the possibility that the relationship between dispersal and polyandry will be dependent on the ecological and demographic context, and on other species’ life-histories. However, we lack theoretical predictions on how dispersal and polyandry may covary under range expansion conditions, under different timing of dispersal relative to mating, as well as we lack empirical estimates of such covariance under both static and dynamic spatial structure. There is therefore large and interesting potential to step-up and expand our knowledge on the evolution of dispersal and mating systems by embracing their joint evolution and feedbacks, theoretically as well as empirically.

## Acknowledgments

I am grateful to J. M. Reid and J. M. J. Travis as this work would not have been possible without their mentorship. I thank A. B. Duthie, M. Tschol, A. Charmouh and L. Dunan for helpful discussions. This work was supported by a Royal Society University Research Fellowship to GB (UF160614), and initially by the European Research Council through funding to J.M. Reid. All simulations were performed on the University of Aberdeen HPC, Maxwell.

## Supplementary Material

**Figure S1.**
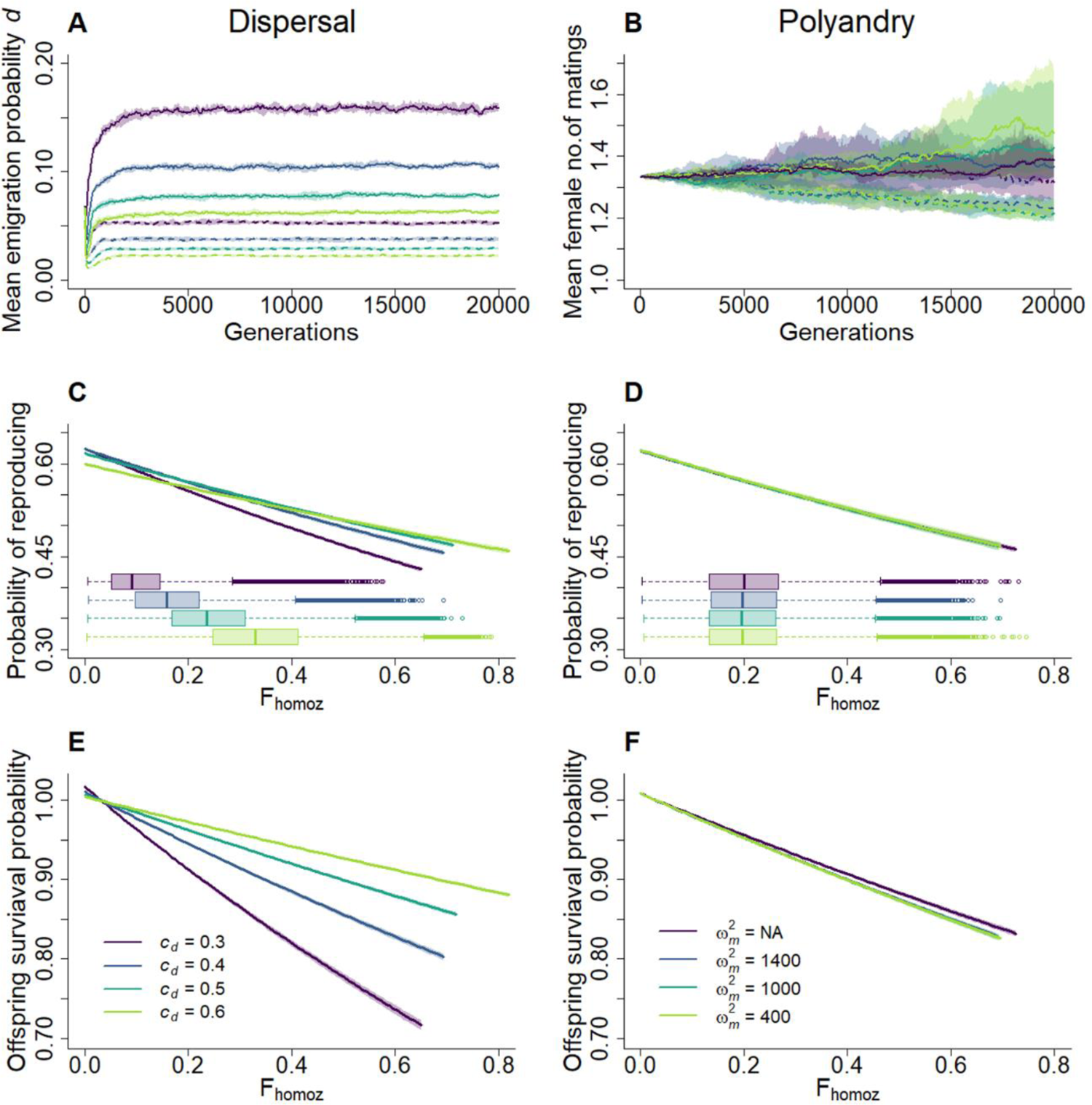
Effect of lower rate of deleterious mutations (*U_d_* = 0.5; *U_l_* = 0.1) on the evolution of dispersal and polyandry when either one or the other trait evolves. A) Evolution of mean dispersal probability phenotypes *d* in the absence of polyandry (*a* = 3.0), under different costs of dispersal (*c_d_* = 0.3, 0.4, 0.5, 0.6), in the absence (dashed lines) or presence (solid lines) of deleterious mutations. **B)** Evolution of mean polyandry phenotypes (expected female number of matings, *P* = 1 + 1/*a*) evolved under fix dispersal probability (*d* = 0.05), as a function of different strengths of direct selection against female remating (no cost; ω_m_^2^ = 1400, 1000, 400) in the absence (dashed lines) or presence (solid lines) of deleterious mutations. In A-B, lines represent the median of mean phenotypes across 20 replicate simulations; colored shades depict the first and third quartile. The color legend for panels A,C,E is presented in E; the legend for panels B,D,F is presented in F. **C-D)** Relationship between individual probability of reproducing and inbreeding coefficient *F_homoz_* (i.e., ID in reproduction probability) when **C)** dispersal evolves under different costs in the absence of polyandry and, **D)** polyandry evolves under different strengths of direct selection with fix dispersal probability. Lines show the fitted models and colored shades the 95% CI. Models are fitted at generation 20,000 to a subsample of 140 populations, across 10 replicates. Boxplots represents the distribution of the individual *F_homoz_*. **E-F)** Relationship between offspring survival probability and *F_homoz_* (i.e., ID in offspring survival probability). In E simulation scenarios and parameters as in C; in F as in D.

**Figure S2.**
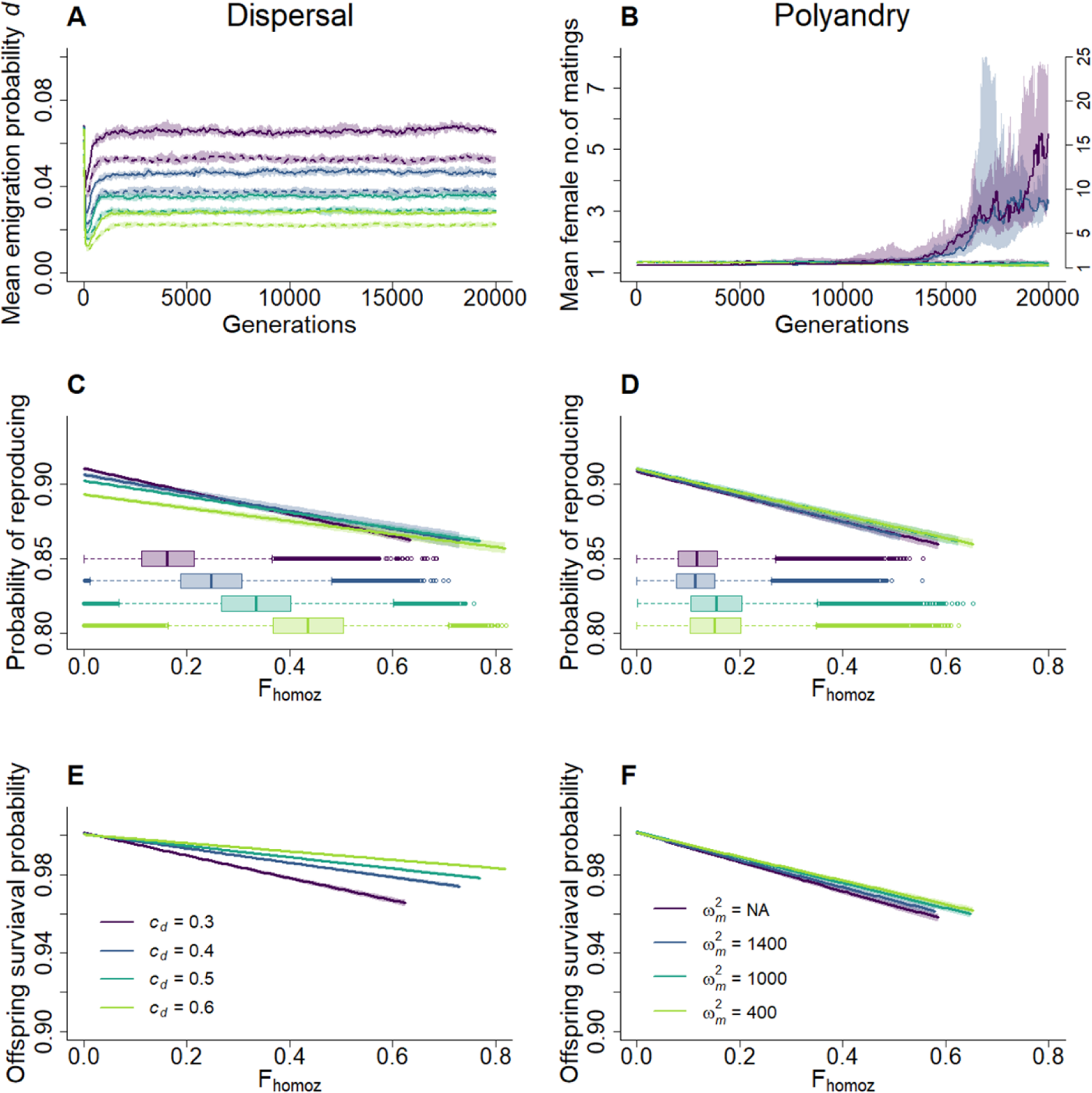
Effect of very low rate of deleterious mutations (*U_d_* = 0.1; *U_l_* = 0.02) on the evolution of dispersal and polyandry when either one or the other trait evolves. A) Evolution of mean dispersal probability phenotypes *d* in the absence of polyandry (*a* = 3.0), under different costs of dispersal (*c_d_* = 0.3, 0.4, 0.5, 0.6), in the absence (dashed lines) or presence (solid lines) of deleterious mutations. B) Evolution of mean polyandry phenotypes (expected female number of matings, *P* = 1 + 1/*a*) evolved under fix dispersal probability (*d* = 0.05), as a function of different strengths of direct selection against female remating (no cost; ω_m_^2^ = 1400, 1000, 400) in the absence (dashed lines) or presence (solid lines) of deleterious mutations. In A-B, lines represent the median of mean phenotypes across 20 replicated simulations; colored shades depict the first and third quartile. The color legend for panels A,C,E is presented in E; the legend for panels B,D,F is presented in F. C-D) Relationship between individual probability of reproducing and inbreeding coefficient *F_homoz_* (i.e., ID in reproduction probability) when C) dispersal evolves under different costs in the absence of polyandry and, D) polyandry evolves under different strengths of direct selection with fix dispersal probability. Lines show the fitted models and colored shades the 95% CI. Models are fitted at generation 20,000 to a subsample of 140 populations, across 10 replicates. Boxplots represents the distribution of the individual *F_homoz_*. E-F) Relationship between offspring survival probability and *F_homoz_* (i.e., ID in offspring survival probability). In E simulation scenarios and parameters as in C; in F as in D.

**Figure S3.**
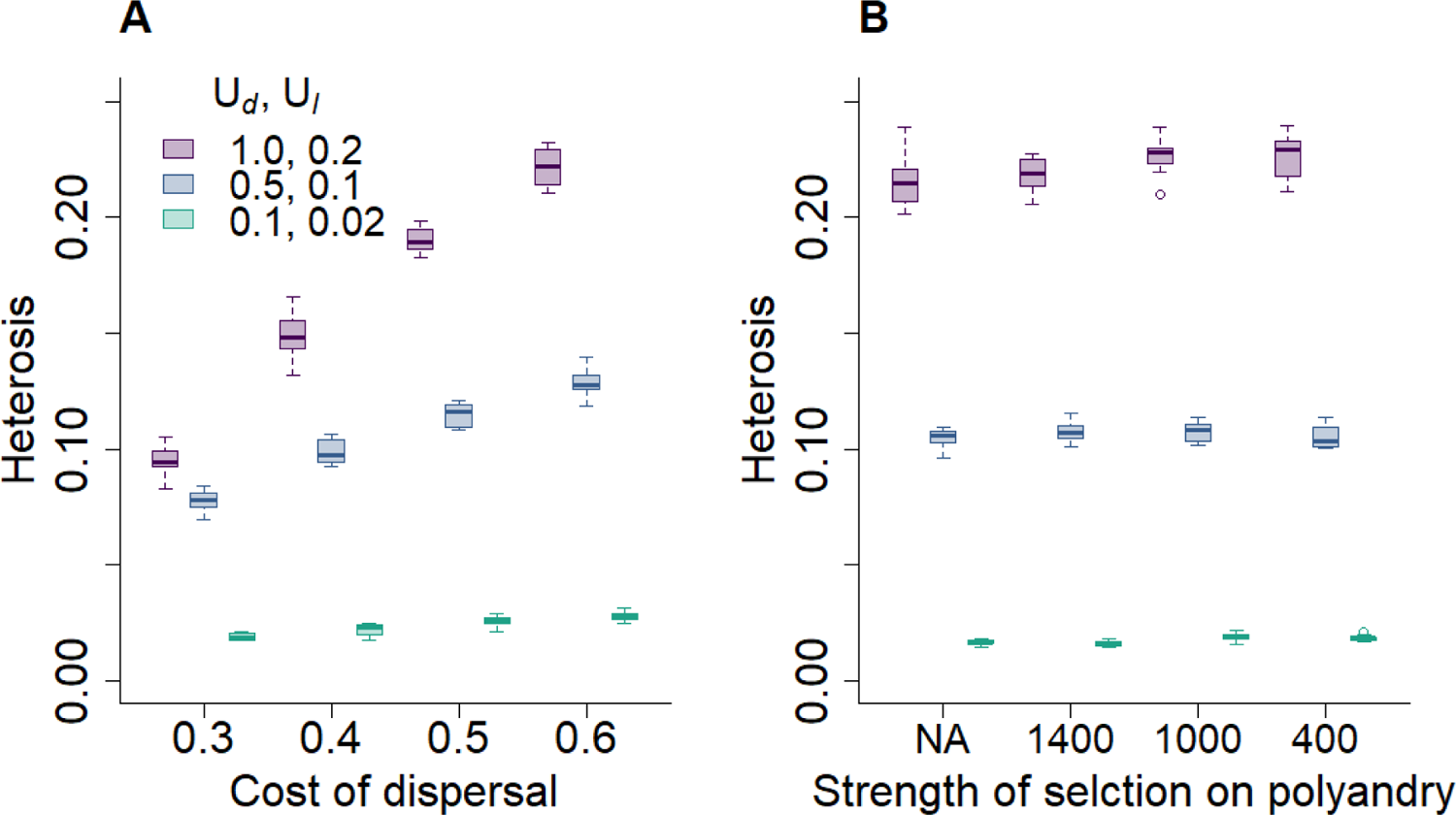
Heterosis emerging when either dispersal or polyandry evolve under different rates of deleterious mutations. A) Heterosis as a function of cost of dispersal *c_d_* and different rates of mildly deleterious (*U_d_*) and lethal (*U_l_*) mutations (colors), when only dispersal is evolving in the absence of polyandry. B) Heterosis as a function of the strength of direct selection against polyandry ω_m_^2^ and different rates deleterious mutations, when only polyandry is evolving under fix dispersal probability (*d* = 0.05). Heterosis is shown as median (solid bands), first and third quartiles (box limits), and approximately twice the standard deviation (whiskers) over 20 replicate simulations at generation 20,000.

**Figure S4.**
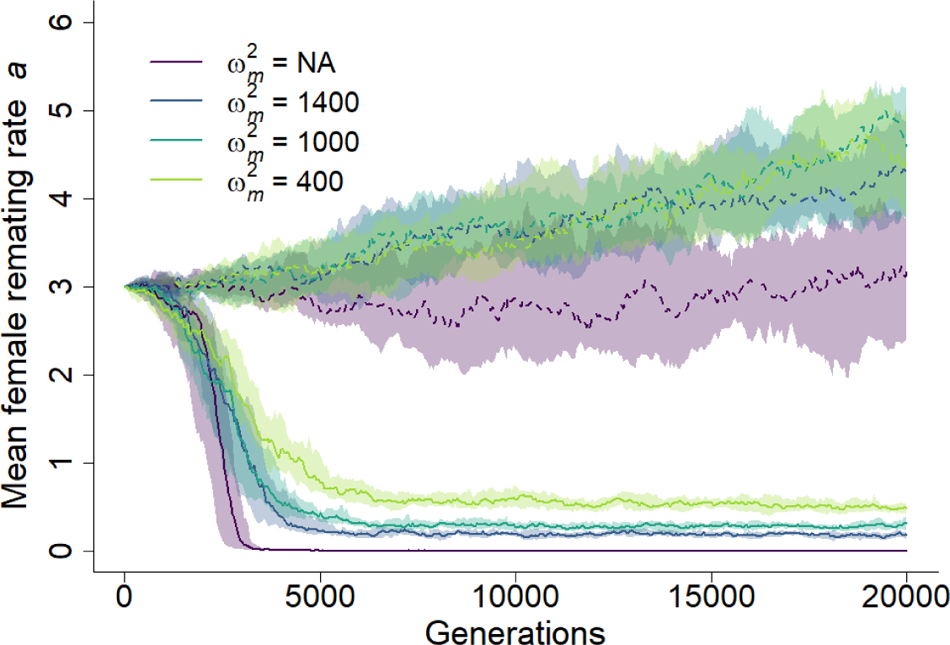
Evolution of female re-mating rate *a* under fix dispersal. Evolution of mean female remating rate phenotypes (*a*) under fix dispersal probability (*d* = 0.05), as a function of different strengths of direct selection against female remating (no cost; ω_m_^2^= 1400, 1000, 400) in the absence (dashed lines) or presence (solid lines) of deleterious mutations. Lines represent the median of mean phenotypes across 20 replicated simulations; colored shades depict the first and third quartile. Other parameters: *U_d_* = 1.0; *U_l_* = 0.2.

**Figure S5.**
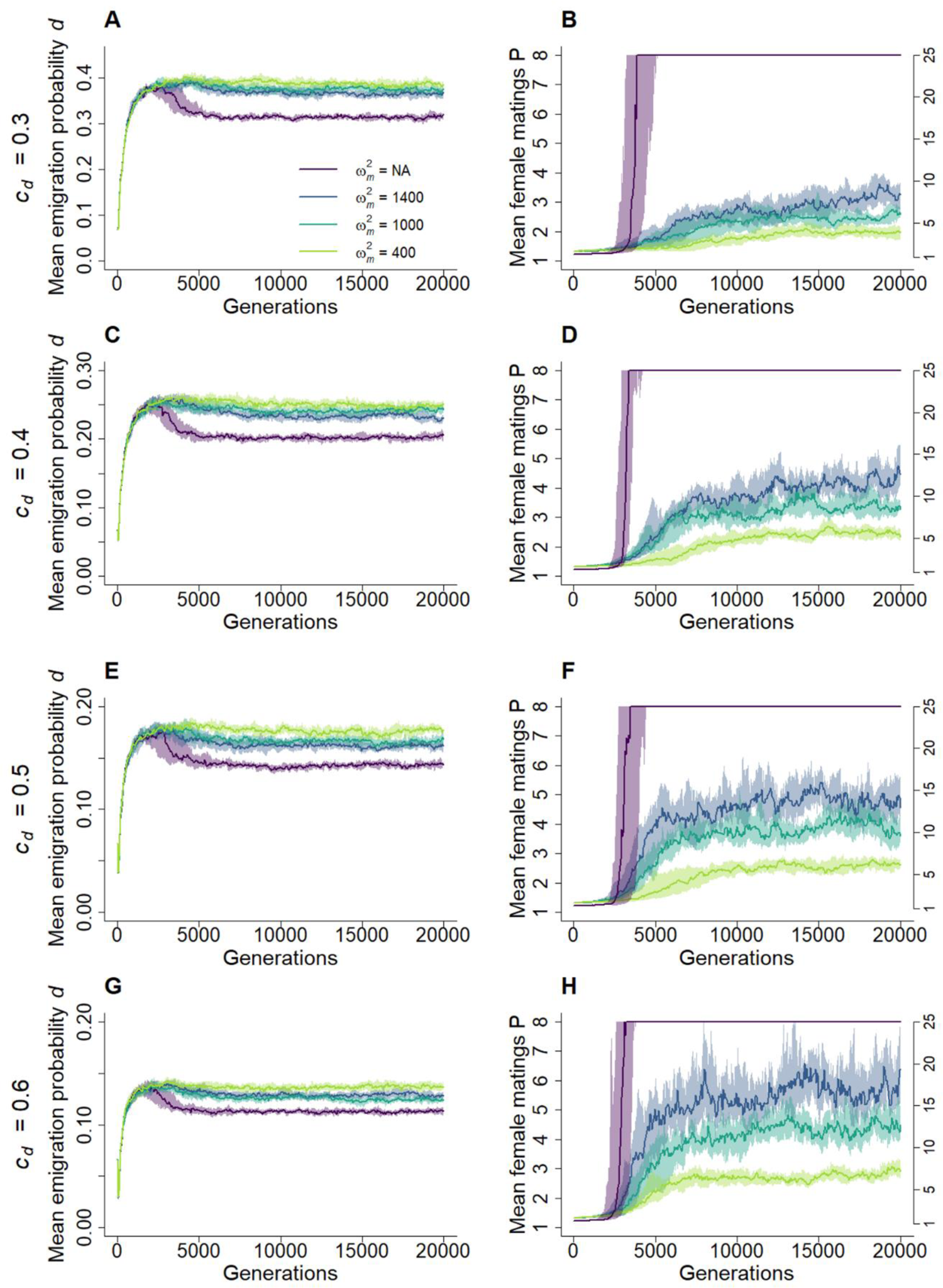
(previous page). Evolutionary dynamics of dispersal and polyandry. Joint evolution of mean dispersal probability phenotypes (*d*) and mean polyandry phenotypes (expect female number of matings: *P* = 1 + 1/*a*) in the presence of deleterious mutation, given different costs of dispersal *c_d_* (**A-B:** 0.3; **C-D:** 0.4; **E-F:** 0.5; **G-H:** 0.6) and different strengths of direct selection against female re-mating (no cost; ω_m_^2^ = 1400, 1000, 400, depicted by different colors). In the absence of direct selection against polyandry (ω_m_^2^= NA), females evolved to mate with all the males in the population; the y-axis on the right hand-side refers to this single line (purple). Lines represent the median of mean phenotypes across 20 replicated simulations; colored shades depict the first and third quartile.

**Figure S6.**
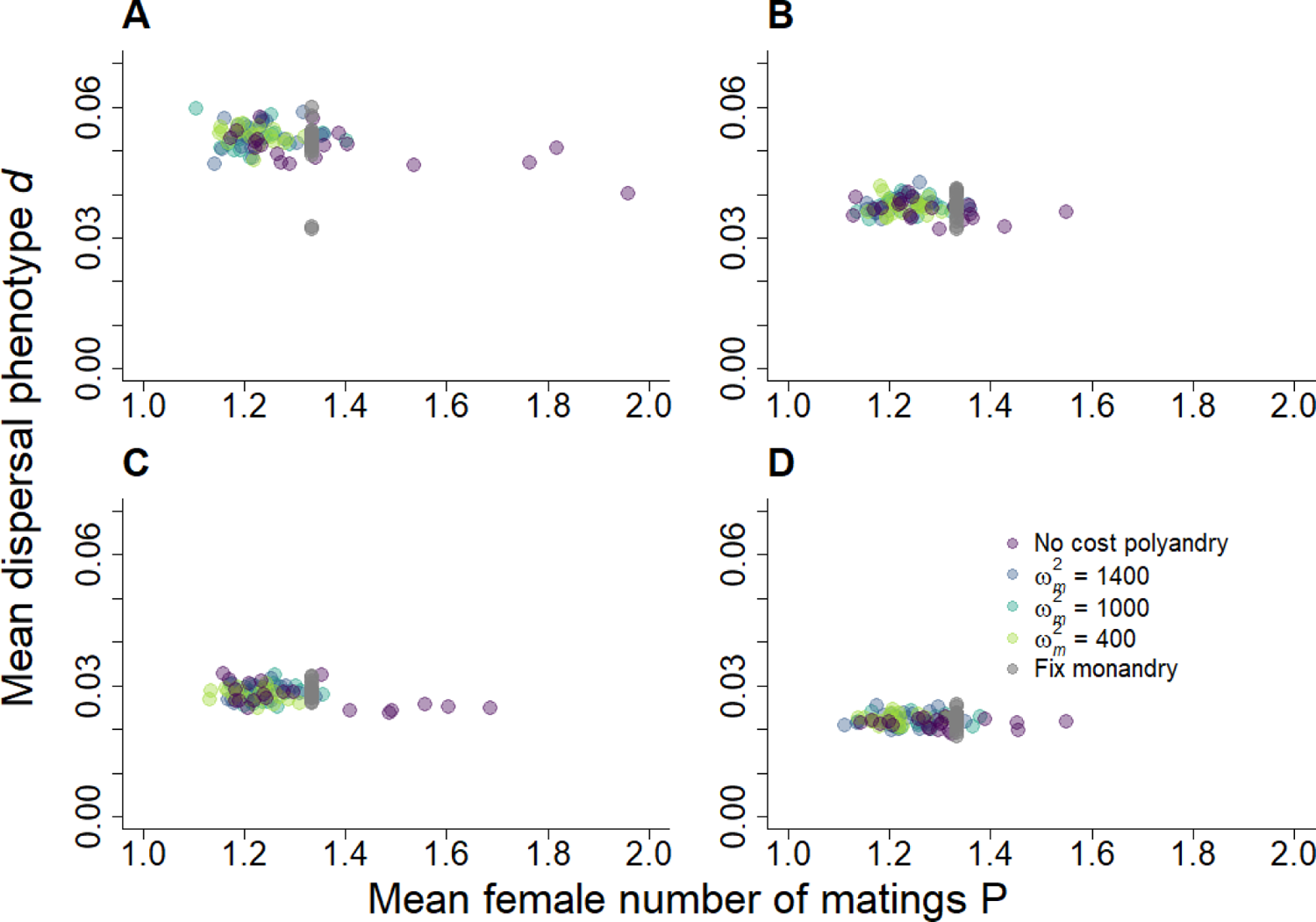
Joint evolution of dispersal and polyandry in the absence of inbreeding depression. Mean dispersal probability phenotypes (*d*) and mean polyandry phenotypes (expected female number of matings: *P* = 1 + 1/*a*) in the absence of deleterious mutations, given different costs of dispersal *c_d_* (**A:** 0.3; **B:** 0.4; **C:** 0.5; **D:** 0.6) and different strengths of direct selection against female re-mating (no cost; ω_m_^2^ = 1400, 1000, 400). Each data point represents the mean phenotypic values for one out of 20 replicate simulations at generation 20,000. Colored dots indicate simulations where dispersal and polyandry jointly evolved; grey dots, simulations where dispersal evolved given fix monandry (*a* = 3.0).

**Figure S7.**
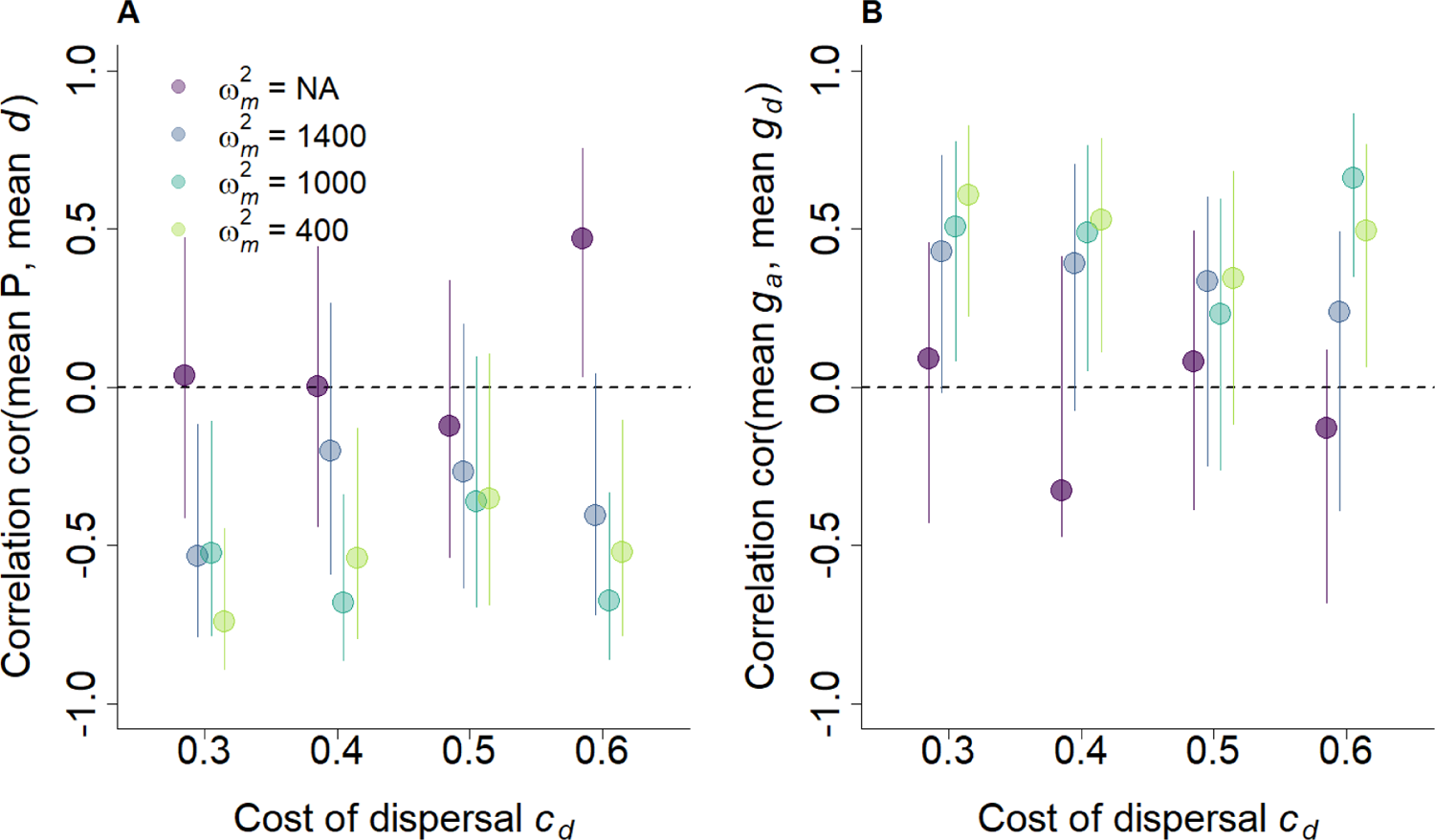
Correlation between polyandry and dispersal at the metapopulation level. **A)** Correlation between mean polyandry phenotype (*P* = 1 + 1/*a*) and mean dispersal phenotype (*d*), and **B)** correlation between mean genotypic female re-mating rate (*g_a_*) and mean genotypic dispersal (*g_d_*), given different costs of dispersal *c_d_* and different strengths of direct selection against female re-mating (no cost; ω_m_^2^ = 1400, 1000, 400). Correlations are calculated across 20 replicate simulations between each replicate mean phenotypic and genotypic values. Means are calculated across all individuals in the metapopulation and over the last 500 generations. Bars represents the 95% confidence intervals.

**Figure S8.**
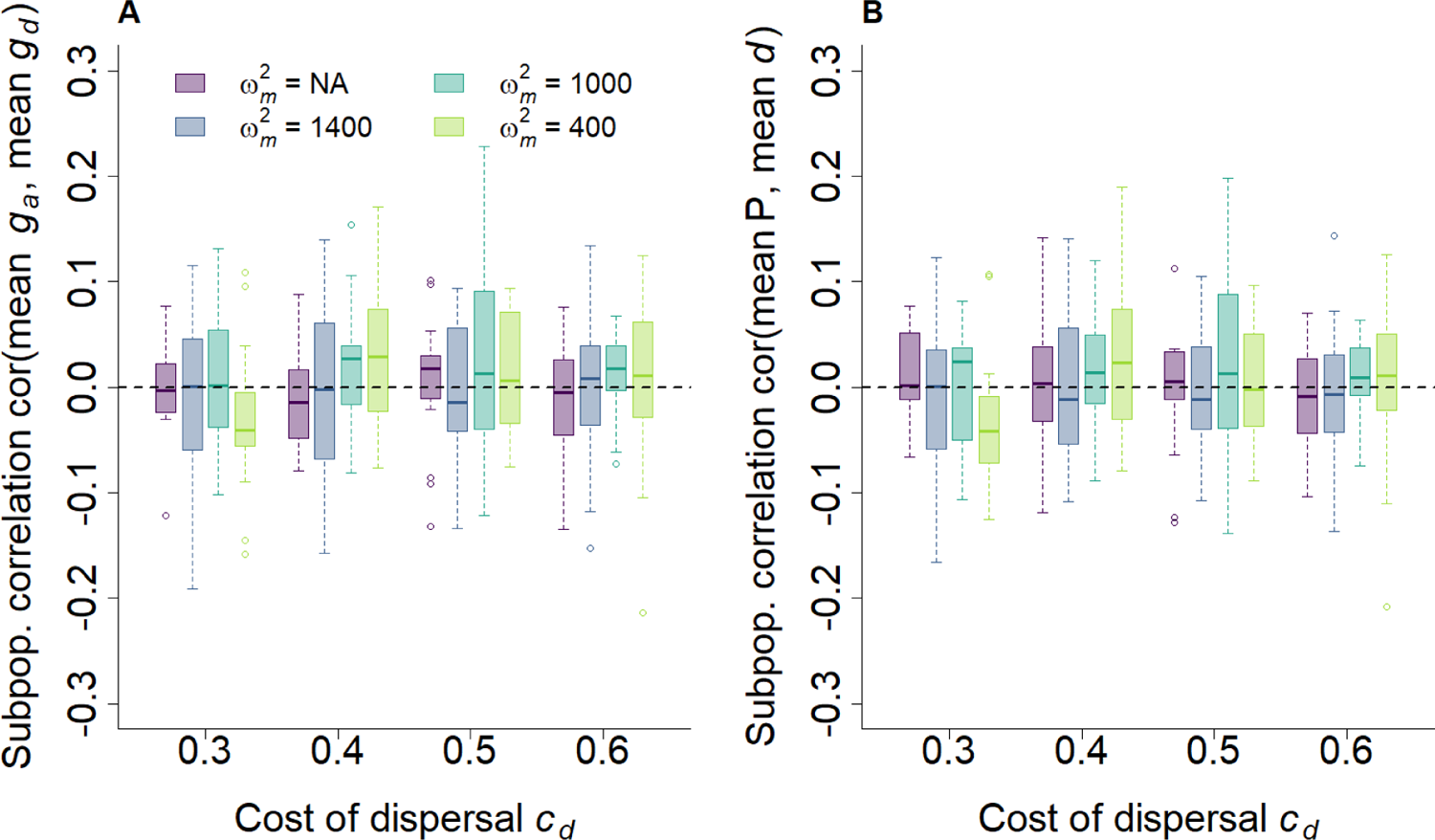
Absence of between subpopulation correlation between polyandry and dispersal. **A)** Correlation between mean genotypic female re-mating rate (*g_a_*) and mean genotypic dispersal (*g_d_*), and **B)** between mean polyandry phenotype (*P* = 1 + 1/*a*) and mean dispersal phenotype (*d*), given different costs of dispersal *c_d_* and different strengths of direct selection against female re-mating (no cost; ω_m_^2^ = 1400, 1000, 400). Correlations are calculated for each replicate across subpopulations mean phenotypic and genotypic values at generation 20,000. Means are calculated across all individuals in each subpopulation. Correlations are shown as median (solid bands), first and third quartiles (box limits), and approximately twice the standard deviation (whiskers) over 20 replicate simulations.

**Figure S9.**
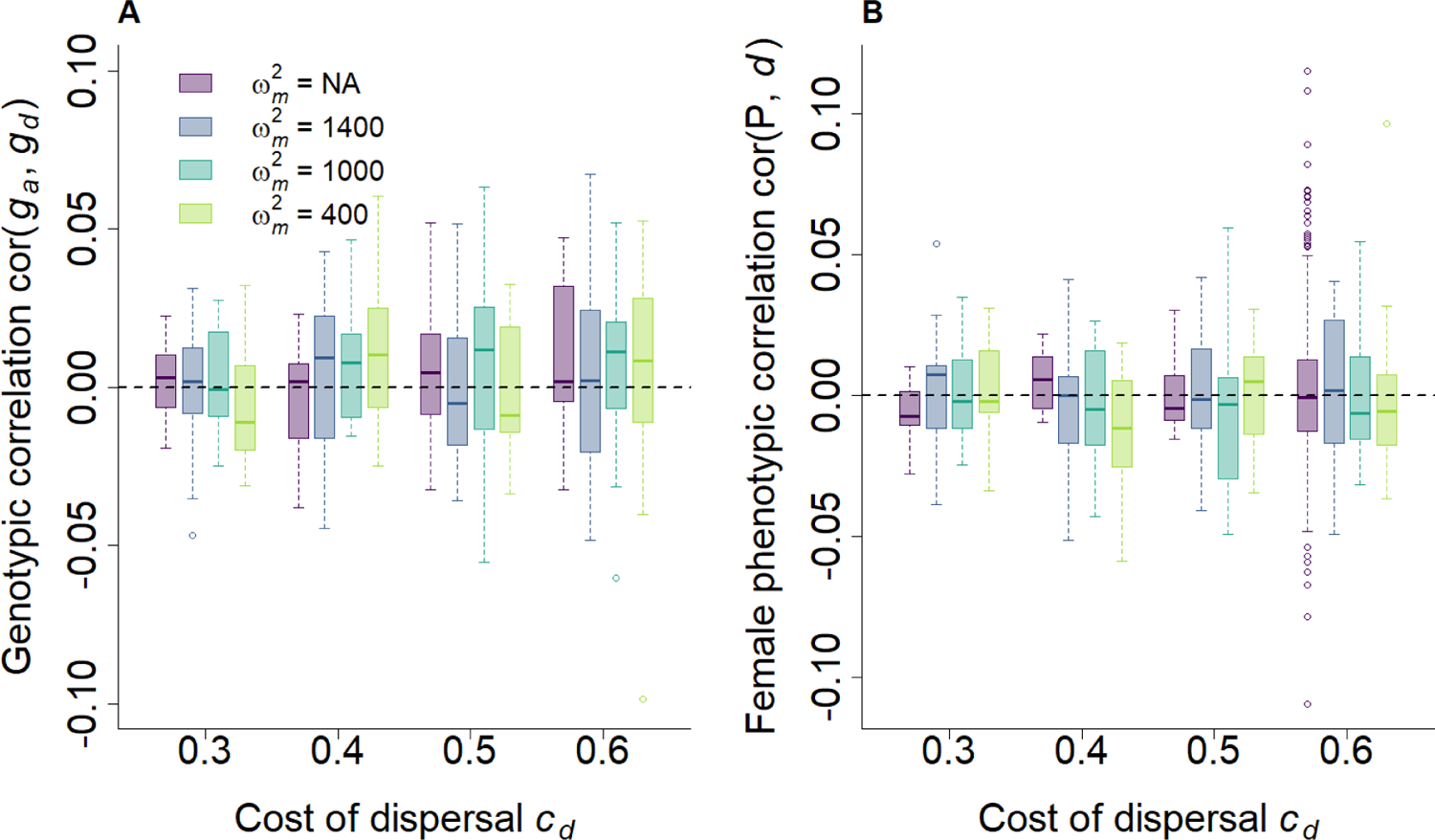
Absence of genetic correlation between polyandry and dispersal. **A)** Genetic correlation between female re-mating rate genotypic value (*g_a_*) and dispersal genotypic value (*g_d_*), and **B)** genetic correlation between female polyandry phenotype (*P* = 1 + 1/*a*) and female dispersal phenotype (*d*), given different costs of dispersal *c_d_* and different strengths of direct selection against female re-mating (no cost; ω_m_^2^ = 1400, 1000, 400). Correlations are calculated for each replicate across individuals phenotypic and genotypic values at generation 20,000, and are shown as median (solid bands), first and third quartiles (box limits), and approximately twice the standard deviation (whiskers) over 20 replicate simulations.

**Figure S10.**
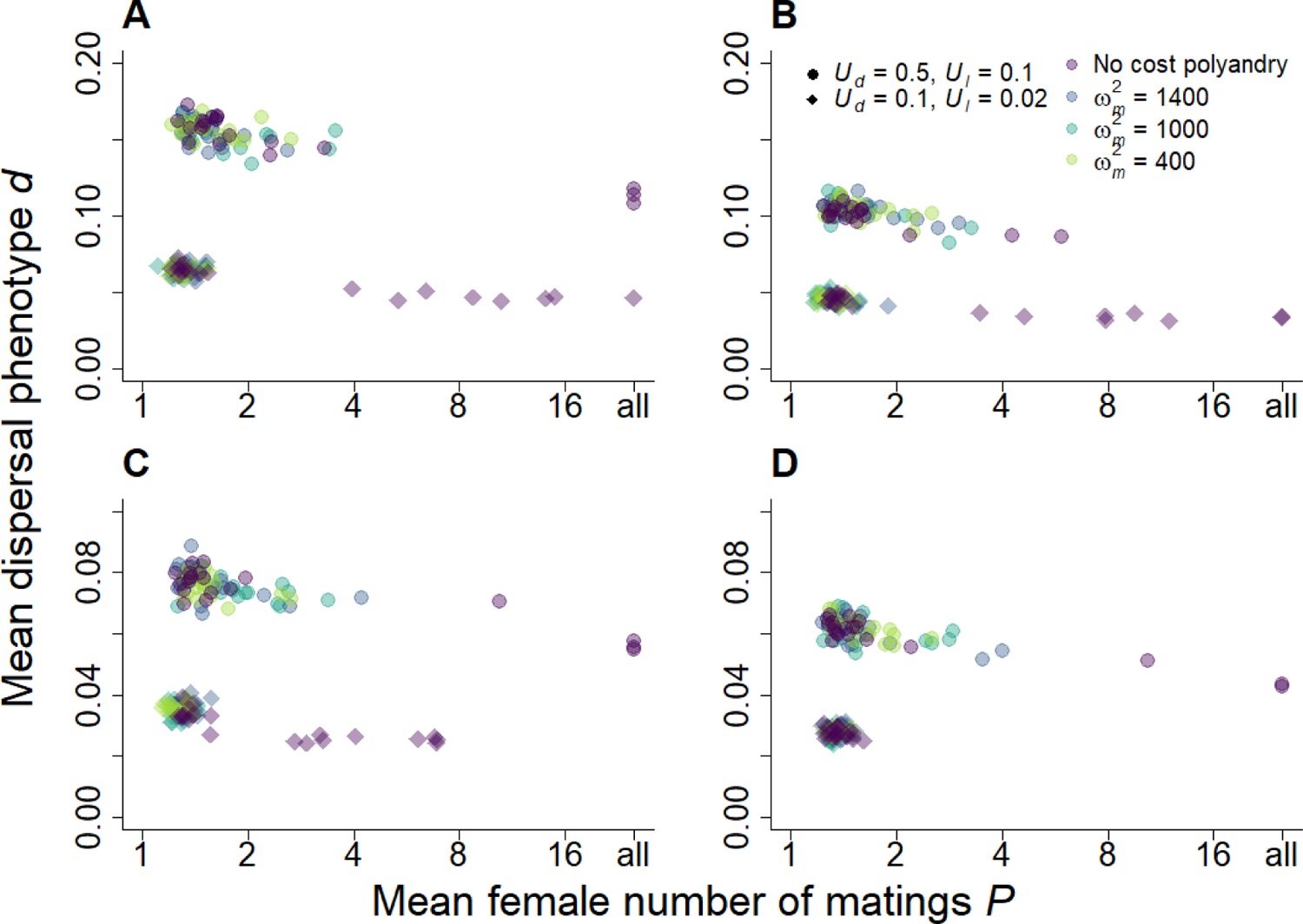
Effect of lower deleterious mutation rate on the joint evolution of dispersal and polyandry. Joint evolution of mean dispersal probability phenotypes (*d*) and mean polyandry phenotypes (expect female number of matings: *P* = 1 + 1/*a*) at two different levels of deleterious mutation rates, given different costs of dispersal *c_d_* (**A:** 0.3; **B:** 0.4; **C:** 0.5; **D:** 0.6) and different strengths of direct selection against female re-mating (no cost; ω^2^_m_= 1400, 1000, 400). Dots represent simulations where *U_d_* = 0.5 and *U_l_* = 0.1, while diamonds represent simulations where *U_d_* = 0.1 and *U_l_* = 0.02, thus corresponding to mutation rates that are half and a tenth, respectively, of the mutation rates presented in the main results (Fig. 2). Each data point represents the mean phenotypic value for one out of 20 replicate simulations at generation 20,000. The x-axis is on the logarithmic scale to aid visualization.

**Figure S11.**
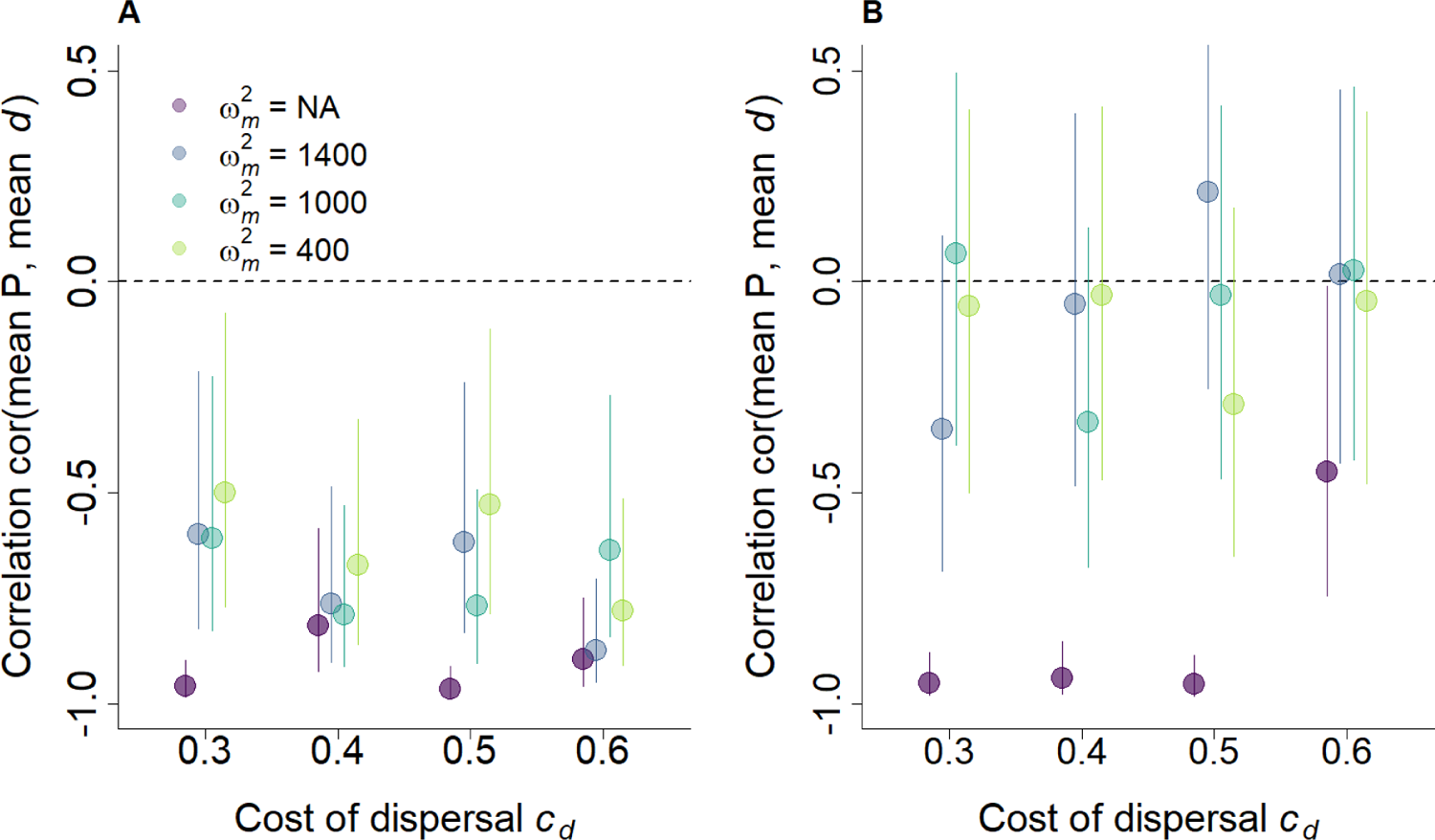
Correlation between polyandry and dispersal at the metapopulation level for lower rates of deleterious mutations. Correlation between mean polyandry phenotype (*P* = 1 + 1/*a*) and mean dispersal phenotype (*d*) given **A)** *U_d_* = 0.5 and *U_l_* = 0.1, and **B)** *U_d_* = 0.1 and *U_l_* = 0.02. Results are presented for different costs of dispersal *c_d_* and different strengths of direct selection against female re-mating (no cost; ω^2^_m_ = 1400, 1000, 400). Correlations are calculated across 20 replicate simulations between each replicate mean phenotypic and genotypic values. Means are calculated across all individuals in the metapopulation and over the last 500 generations. Bars represents the 95% confidence intervals.

**Figure S12.**
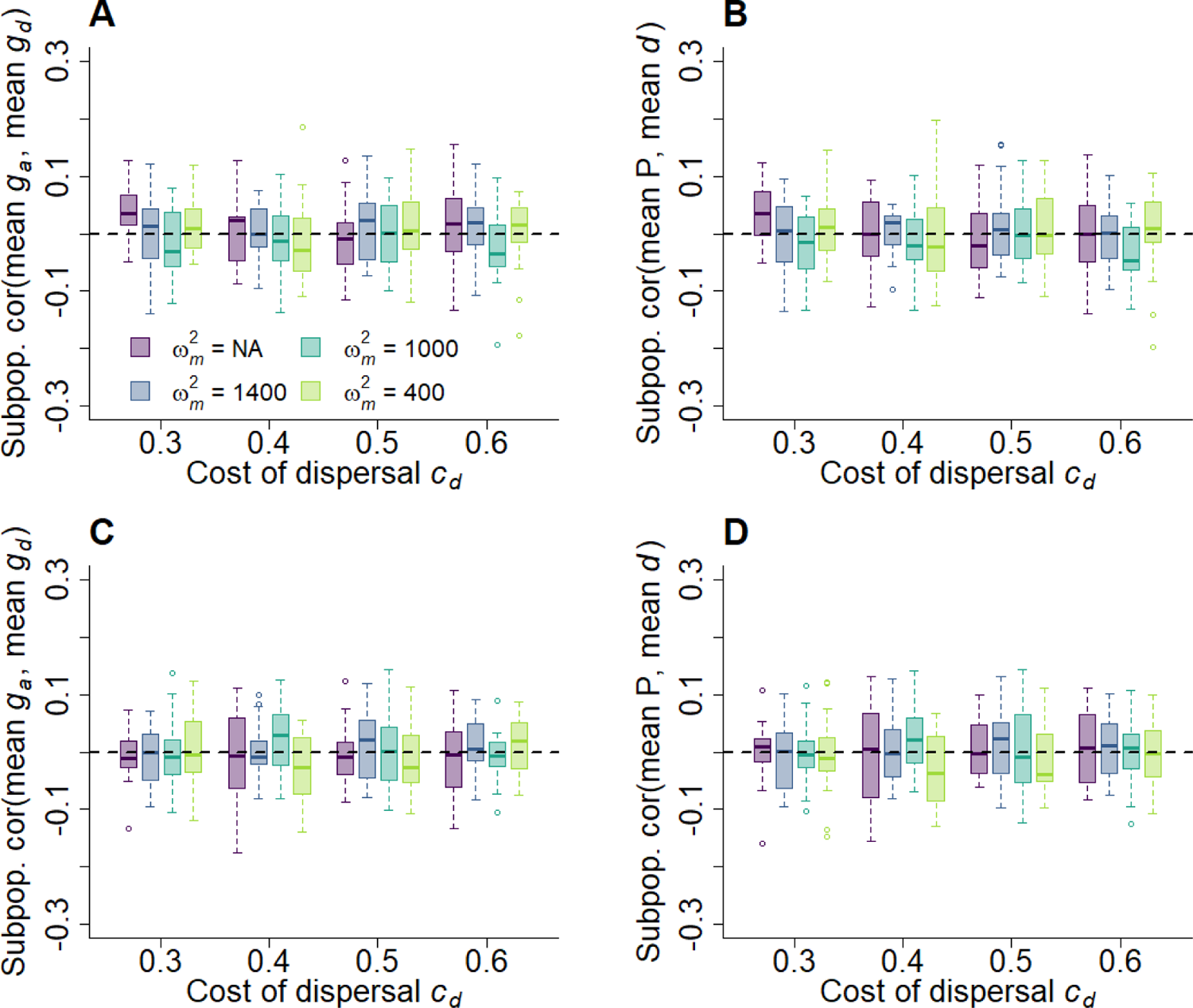
Correlation between polyandry and dispersal at the subpopulation level for lower rates of deleterious mutations. **A)** Correlation between mean genotypic female re-mating rate (*g_a_*) and mean genotypic dispersal (*g_d_*), and **B)** between mean polyandry phenotype (*P* = 1 + 1/*a*) and mean dispersal phenotype (*d*), given different costs of dispersal *c_d_* and different strengths of direct selection against female re-mating (no cost; ω^2^_m_ = 1400, 1000, 400), when mutation rates *U_d_* = 0.5 and *U_l_* = 0.1. **C-D)** same as A-B, but when *U_d_* = 0.1 and *U_l_* = 0.02. Correlations are calculated for each replicate across subpopulations mean phenotypic and genotypic values at generation 20,000. Means are calculated across all individuals in each subpopulation. Correlations are shown as median (solid bands), first and third quartiles (box limits), and approximately twice the standard deviation (whiskers) over 20 replicate simulations.

**Figure S13.**
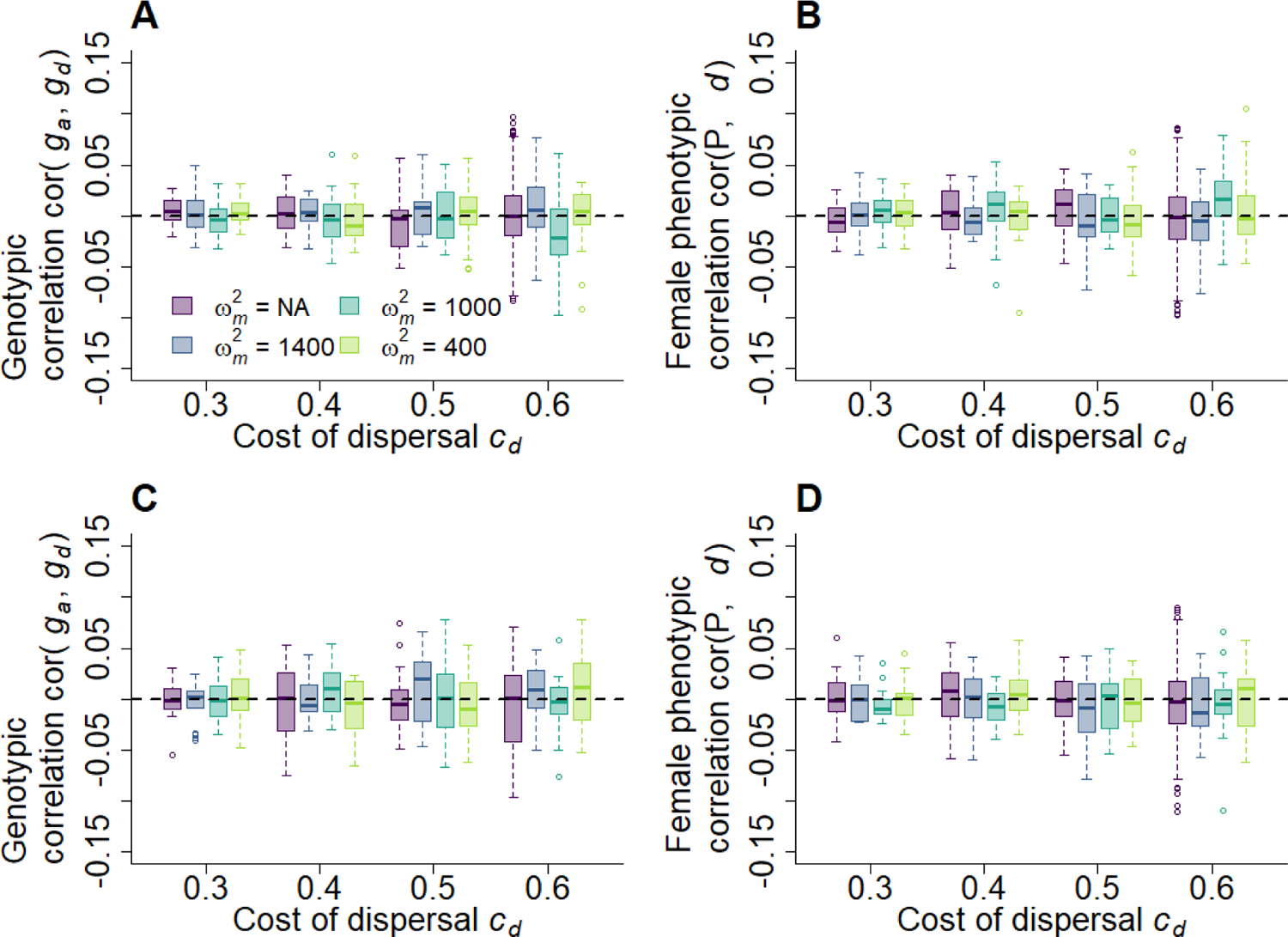
Genetic correlation between polyandry and dispersal for lower rates of deleterious mutations. **A)** Genetic correlation between female re-mating rate genotypic value (*g_a_*) and dispersal genotypic value (*g_d_*), and **B)** genetic correlation between female polyandry phenotype (*P* = 1 + 1/*a*) and female dispersal phenotype (*d*), given different costs of dispersal *c_d_* and different strengths of direct selection against female re-mating (no cost; ω^2^_m_ = 1400, 1000, 400), when mutation rates *U_d_* = 0.5 and *U_l_* = 0.1. **C-D)** same as A-B, but when *U_d_* = 0.1 and *U_l_* = 0.02. Correlations are calculated for each replicate across individuals phenotypic and genotypic values at generation 20,000, and are shown as median (solid bands), first and third quartiles (box limits), and approximately twice the standard deviation (whiskers) over 20 replicate simulations.

---

**Table S1.** Model variables and parameters.

| Variables | Description | Parameter values |
| --- | --- | --- |
| $K$ | Carrying capacity per cell (sub-population) | 50 individuals |
| $f$ | Female fecundity | 12 |
| **Inbreeding and inbreeding depression** |  |  |
| $R$ | Genome map length (for the continuous chromosome carrying deleterious mutations) | 10 |
| $s$ | Selection coefficient of mildly deleterious mutations | $\Gamma(\alpha, s_d/\alpha)$ |
| $h$ | Dominance coefficient of mildly deleterious mutations | $U[0.0, e^{-ks}]$ |
| $s_d$ | Mean selection coefficient of mildly deleterious mutations | 0.05 |
| $h_d$ | Mean dominance coefficient of mildly deleterious mutations | 0.3 |
| $\alpha$ | Shape parameter of gamma distribution of $s$ | 1.0 |
| $k$ | Constant | $-\log(2h_d)/s_d$ |
| $s_l$ | Selection coefficient of lethal mutations | 1.0 |
| $h_l$ | Dominance coefficient of lethal mutations | 0.02 |
| $U_d$ | Mutation rate for mildly deleterious mutations | $1^1, 0.5, 0.1$ / diploid genome / generation |
| $U_l$ | Mutation rate for lethal mutations | $0.2^*, 0.1, 0.02$ / diploid genome / generation |
| $L_n$ | Number of neutral diploid loci | 500 |
| | Neutral allelic values | $U[-1000.0, 1000.0]$ |
| $F_{\text{homoz}}$ | Individual neutral homozygosity (proxy for inbreeding coefficient) | |
| $r$ | Recombination rate for neutral loci | 0.1 |
| $\mu_n$ | Mutation probability for neutral alleles | 0.001 / haploid locus / generation |
| **Traits** |  |  |
| $L$ | Number of diploid loci for each trait | 1 |
| $a$ | Female re-mating interval (phenotypic value) | $a \geq 0.0$ |
| $d$ | Emigration probability (phenotypic value) | $0.0 \leq d \leq 1.0$ |
|  | Initial genotypic mean for female re-mating rate | 3.0 |

<sup>1</sup> Values used in the simulations presented in the main text.

|  |  |  |
| --- | --- | --- |
|  | Initial genotypic mean for emigration probability | 0.05 |
| $\sigma^2_{a,0}$ | Initial genotypic variance for female re-mating rate | 0.25 |
| $\sigma^2_{d,0}$ | Initial genotypic variance for emigration probability | 0.1 |
| $\mu$ | Mutation probability | 0.001 / haploid locus / generation |
| | Mutational effects | $N(0.0, 0.07)$ |
| **Costs** |  |  |
| $\omega^2_m$ | Strength of direct selection against female multiple mating | no cost, 1400, 1000, 400 |
| $c_d$ | Cost of dispersal | 0.3, 0.4, 0.5, 0.6 |

**Table S2.** Coefficients (i.e., mutation load *β_0_*, and inbreeding load *β_1_*) and relative standard errors of the relationship between individual probability of reproducing and inbreeding coefficient *F_homoz_*, and between offspring survival probability and *F_homoz_*. Results are presented for simulations where dispersal evolves under different costs (*c_d_*), in the absence of polyandry, and where polyandry evolves under different strengths of direct selection (ω_m_^2^), with fix dispersal probability. Models are fitted at generation 20,000 to a subsample of 140 populations, across 10 replicates.

| Probability of reproducing |  |  |  |  | Offspring survival |  |  |  |
| --- | --- | --- | --- | --- | --- | --- | --- | --- |
| Only dispersal evolving |  |  |  |  |  |  |  |  |
| $C_d$ | $\beta_0$ | SE | $\beta_1$ | SE | $\beta_0$ | SE | $\beta_1$ | SE |
| 0.3 | -0.99 | 0.005 | -1.34 | 0.016 | 0.11 | 0.001 | -1.97 | 0.025 |
| 0.4 | -0.94 | 0.003 | -1.16 | 0.018 | 0.07 | 0.001 | -1.29 | 0.014 |
| 0.5 | -0.92 | 0.005 | -0.98 | 0.016 | 0.04 | 0.001 | -0.78 | 0.008 |
| 0.6 | -0.92 | 0.002 | -0.79 | 0.014 | 0.02 | 0.0004 | -0.52 | 0.005 |
| Only polyandry evolving |  |  |  |  |  |  |  |  |
| $\omega_m^2$ | $\beta_0$ | SE | $\beta_1$ | SE | $\beta_0$ | SE | $\beta_1$ | SE |
| none | -0.94 | 0.004 | -0.81 | 0.007 | 0.02 | 0.0003 | -0.49 | 0.005 |
| 1400 | -0.94 | 0.003 | -0.79 | 0.014 | 0.02 | 0.0003 | -0.46 | 0.005 |
| 1000 | -0.94 | 0.003 | -0.78 | 0.009 | 0.02 | 0.0003 | -0.46 | 0.005 |
| 400 | -0.94 | 0.003 | -0.77 | 0.009 | 0.02 | 0.0003 | -0.46 | 0.005 |

**Table S3.** Jointly evolving dispersal and polyandry affect evolution of inbreeding depression in reproduction and survival. Coefficients (i.e., mutation load *β_0_*, and inbreeding load *β_1_*) and relative standard errors of the relationship between individual probability of reproducing and inbreeding coefficient *F_homoz_*, and between offspring survival probability and *F_homoz_*, when dispersal and polyandry are jointly evolving. Results are presented for varying costs of dispersal (*c_d_*) and strengths of direct selection against female multiple mating (ω_m_^2^). Models are fitted at generation 20,000 to a subsample of 140 populations, across 10 replicates.

| $C_d$ | $\omega_m^2$ | *Probability of reproduction* | | | | *Survival probability* | | | |
| --- | --- | --- | --- | --- | --- | --- | --- | --- | --- |
| | | $\beta_0$ | SE | $\beta_1$ | SE | $\beta_0$ | SE | $\beta_1$ | SE |
| **0.3** | **none** | -0.99 | 0.003 | -1.29 | 0.024 | 0.07 | 0.001 | -1.74 | 0.023 |
|  | **1400** | -0.99 | 0.002 | -1.35 | 0.022 | 0.09 | 0.001 | -1.90 | 0.024 |
|  | **1000** | -0.99 | 0.003 | -1.34 | 0.020 | 0.09 | 0.001 | -1.86 | 0.024 |
|  | **400** | -0.99 | 0.004 | -1.35 | 0.017 | 0.09 | 0.001 | -1.87 | 0.024 |
| **0.4** | **none** | -0.95 | 0.003 | -1.09 | 0.035 | 0.04 | 0.001 | -1.10 | 0.013 |
|  | **1400** | -0.95 | 0.001 | -1.14 | 0.016 | 0.05 | 0.001 | -1.20 | 0.014 |
|  | **1000** | -0.95 | 0.004 | -1.17 | 0.014 | 0.05 | 0.001 | -1.22 | 0.014 |
|  | **400** | -0.95 | 0.002 | -1.17 | 0.019 | 0.06 | 0.001 | -1.20 | 0.014 |
| **0.5** | **none** | -0.94 | 0.003 | -0.93 | 0.014 | 0.03 | 0.0004 | -0.72 | 0.008 |
|  | **1400** | -0.93 | 0.002 | -0.96 | 0.018 | 0.03 | 0.0004 | -0.76 | 0.008 |
|  | **1000** | -0.93 | 0.003 | -0.91 | 0.013 | 0.03 | 0.0004 | -0.74 | 0.008 |
|  | **400** | -0.93 | 0.003 | -0.99 | 0.011 | 0.04 | 0.0005 | -0.78 | 0.008 |
| **0.6** | **none** | -0.94 | 0.002 | -0.77 | 0.012 | 0.02 | 0.0003 | -0.46 | 0.005 |
|  | **1400** | -0.93 | 0.001 | -0.80 | 0.012 | 0.02 | 0.0003 | -0.50 | 0.005 |
|  | **1000** | -0.94 | 0.003 | -0.80 | 0.012 | 0.02 | 0.0003 | -0.49 | 0.005 |
|  | **400** | -0.93 | 0.004 | -0.80 | 0.011 | 0.03 | 0.0003 | -0.52 | 0.006 |

